# *convertible*CARs: A chimeric antigen receptor system for flexible control of activity and antigen targeting

**DOI:** 10.1101/696401

**Authors:** Kyle E. Landgraf, Steven R. Williams, Daniel Steiger, Dana Gebhart, Stephen Lok, David W. Martin, Kole T. Roybal, Kaman Chan Kim

## Abstract

We have developed a chimeric antigen receptor (CAR) platform that functions as a modular system to address limitations of current CAR therapies. An inert form of the NKG2D extracellular domain (iNKG2D) was used as the ectodomain of the CAR to generate *convertible*CAR™-T cells. These cells were activated only when an immunological synapse was formed with an antigenic target, mediated by a bispecific adaptor comprised of an iNKG2D-exclusive ULBP2-based ligand fused to an antigen-targeting antibody (MicAbody^TM^). Efficacy against Raji tumors in NSG mice was dependent upon doses of both a rituximab-based MicAbody and *convertible*CAR-T cells. We have also demonstrated that the exclusive ligand-receptor partnering enabled the targeted delivery of a mutant form of IL-2 to exclusively promote the expansion of *convertible*CAR-T cells *in vitro* and *in vivo*. By altering the Fv domains of the MicAbody or the payload fused to the orthogonal ligand, *convertible*CAR-T cells can be readily targeted or regulated.

## INTRODUCTION

The engineering of patient-derived T cells to express chimeric antigen receptors (CARs) has altered the landscape of adoptive cell therapies, providing scientists and clinicians the ability to harness the powerful cytolytic capabilities of T cells and direct them to specific antigen-expressing targets in an MHC-independent manner. Their initial application to treat hematologic malignancies has led to astounding responses and inspired a surge of research efforts to improve upon the original designs in order to strike the best balance of CAR-T cell potency, safety, and persistence and to drive their effective use in non-hematologic indications^1 2^. Sufficient pre-clinical and clinical data have accumulated from the variations on prototype CAR architecture such that there is now a deeper appreciation of the subtleties that affect CAR-T function. However, CAR-T cell therapies are significantly limited by their utilization of a single-purpose targeting domain, lack of dose control which can contribute to cytokine release syndrome, and inability to address tumor antigen loss leading to disease relapse^3 4^. There is a need to develop a modular CAR-T cell system that enables sequential and multiplex targeting and that, ideally, would provide elements of dose controlled activity, expansion, and persistence. A number of creative solutions have been explored utilizing adaptor molecules that can engage a constant CAR for differential antigen targeting (reviewed in^5^). However, there is no single bio-orthogonal human system that has been demonstrated to have the potential for all the following: flexible targeting to direct T cell activity to antigen of choice, multiplex capabilities to reduce the potential for antigen-loss related relapse, dose control for differential engagement of CAR-T cells, and selective delivery of modulatory agents to CAR-expressing cells.

NKG2D is an activating receptor expressed as a type II homodimeric integral membrane protein on Natural Killer (NK) cells, some myeloid cells and certain T cells^6 7 8^. Human NKG2D has eight distinct natural MIC ligands (MICA, MICB, ULBP1 through ULBP6) that are upregulated on the surface of cells in response to a variety of stresses and their differential regulation provides the immune system a means of responding to a broad range of emergency cues with minimal collateral damage^9 10 11^. The structure of the NKG2D ectodomain, several soluble ligands, and the bound complex of ligands to the ectodomain have been solved, revealing a saddle-like groove in the homodimer interface which engages the structurally conserved α_1_-α_2_ domains of the ligands that are otherwise of disparate amino acid identity^12 13 14 15^. Based upon this structure-function information, we have mutated the NKG2D ectodomain to render it incapable of engaging any natural ligands^16^. Orthogonal α_1_-α_2_ domain ligand variants were then selected to exclusively engage the inert NKG2D (iNKG2D) but not wild-type NKG2D (wtNKG2D). iNKG2D serves as the extracellular component of our CAR construct while the orthogonal U2S3 variant is expressed as a fusion to antigen-specific antibodies, generating bispecifics termed MicAbodies. MicAbodies are capable of directing and activating iNKG2D-CAR-expressing T cells only when the appropriate antigen is displayed on a surface. Keeping the iNKG2D-CAR receptor constant, we have demonstrated that with either single or multiplexed MicAbodies, *convertible*CAR-T cells can be targeted to different tumor antigens to mediate cytolysis. In a disseminated *in vivo* Raji tumor model, efficacy required both iNKG2D-CAR T cells and MicAbody, both in a dose-dependent manner. Lastly, we have determined that U2S3 can be used as a means of addressing molecules for exclusive delivery to iNKG2D-expressing cells and that this feature can specifically target the iNKG2D-CAR cells for complement-mediated killing or drive their expansion *in vitro* and *in vivo* with a mutant cytokine fusion. This highly modular *convertible*CAR system leverages the significant body of information behind antibody discovery, development, and manufacturing and nimbly integrates it with the tremendous potential of adoptive cell therapies.

## RESULTS

### Engineering a privileged binding interaction between an inert NKG2D receptor and orthogonal ULBP2 variant

Two central tyrosine residues in each NKG2D monomer have critical roles in driving receptor-ligand interactions^16^ (Figure 1A). Mutations at these residues were heavily explored, with the Y152A mutant (“iNKG2D.YA”) and the Y152A/Y199F double mutant (“iNKG2D.AF”) selected for further study since both readily expressed as recombinant Fc-fused proteins (data not shown) and were confirmed by biolayer interferometry (BLI) (Supplementary Figs. 1A, 3A) and ELISA (SFig. 1B) to have lost binding to all naturally occurring human ligands. The ULBP2 α_1_α_2_ domain was chosen for phage display-based selection of mutants with high affinity binding to each of the iNKG2D variants since it is not polymorphic^17^. NNK libraries interrogating helix 2 and helix 4 (Figs. 1B, 1C)^14^ returned only helix 4 variants and even then only in the context of a spontaneous R81W mutation which likely has a stabilizing role on the ULBP2 α_1_α_2_ domain. Competitive selection with rounds of increasing concentration of wtNKG2D (SFig. 2A) yielded three variants – U2S1, U2S2, and U2S3 – that reproducibly bound exclusively to iNKG2D.YA even when reformatted as fusions to the C-terminus of the IgG1 heavy chain of the anti-FGFR3 antibody clone R3Mab (SFig. 2B). Although the R81W mutation alone enhanced affinity towards both wtNKG2D and iNKG2D.YA, (SFig. 2B, 2C), its presence in the iNKG2D-selective variants was deemed essential since its reversion to the wild-type residue resulted in loss of binding to iNKG2D.YA (data not shown). As U2S3 consistently exhibited a greater binding differential, it was characterized more thoroughly and shown as a monomer to have a 10-fold higher affinity towards iNKG2D.YA than wild-type ULBP2 had to wtNKG2D (SFig. 2C). Picomolar binding to iNKG2D.YA was measured with a bivalent rituximab antibody fusion and orthogonality was retained by both light-chain (LC) and heavy-chain (HC) fusion configurations (Fig. 1D).

**Figure 1:**
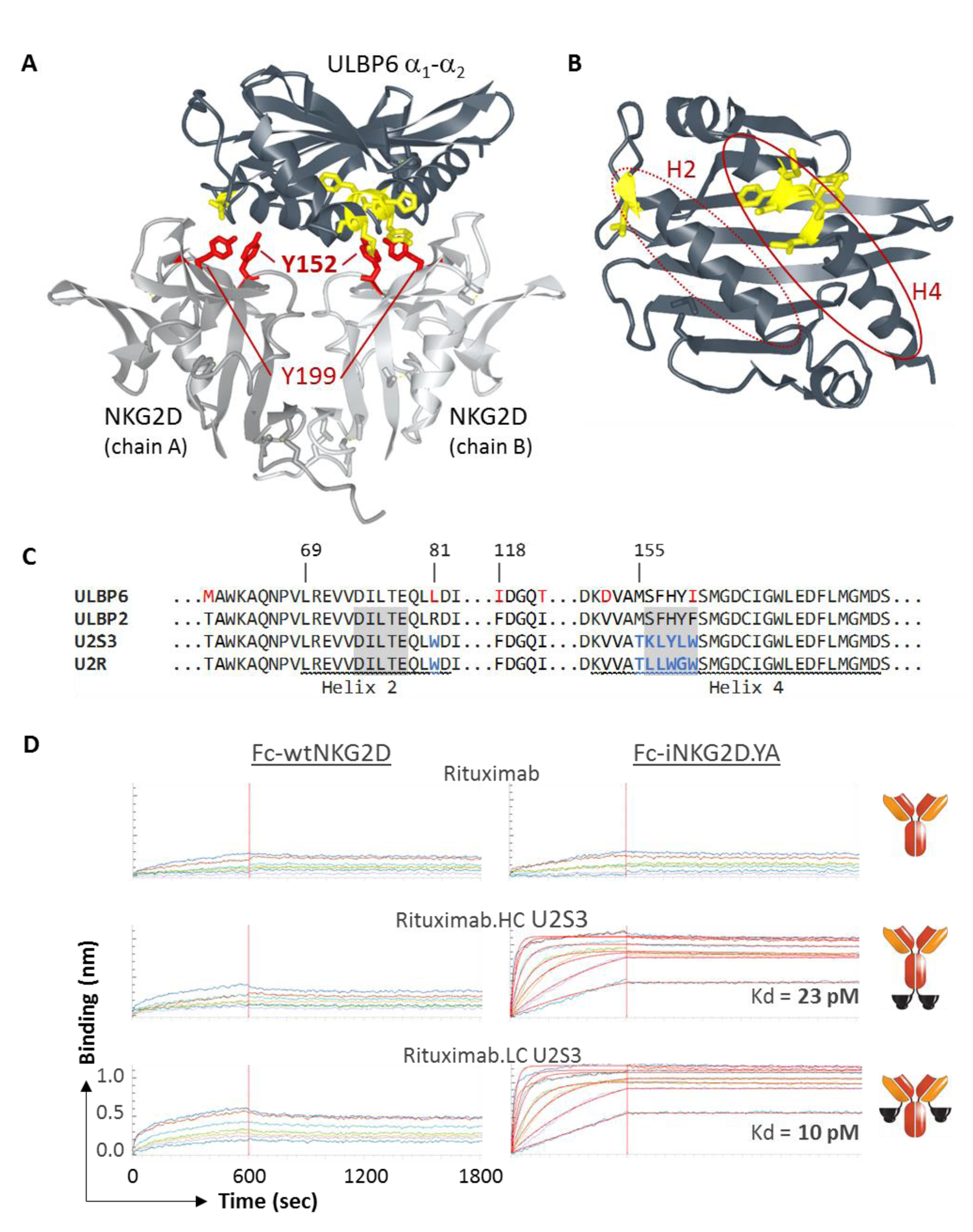
**(A)** The structure of NKG2D in complex with ULBP6 (PDB: 4S0U) was used to illustrate the residues in both the NKG2D homodimer and ligand that were targeted for mutagenesis. Light gray chains correspond to the two NKG2D monomers with Tyrosine 152 and Tyrosine 199, important for ligand engagement, highlighted in red. The dark gray chain corresponds to ULBP6 with the locations of residues 81 and 156-160, mutated in the orthogonal ligands, highlighted in yellow. PDB files were manipulated using the NCBI iCn3D 3D Structure Viewer (https://www.ncbi.nlm.nih.gov/Structure/icn3d/full.html) **(B)** The same ULBP6 structure but with NKG2D absent and the ligand rotated 90° to reveal the two critical helix domains – H2 and H4 (in red ovals) – that engage with the NKG2D homodimer. **(C)** The ULBP2 and ULBP6 α_1_-α_2_ domains are 96.7% identical with the differences in red font. The locations of helices 2 and 4 are underlined, the NNK library residues highlighted in gray, and the residues incorporated into the final orthogonal ligands - ULBP2.S3 (U2S3) and ULBP2.R (U2R) for iNKG2D.YA and iNKG2D.AF, respectively - indicated in blue. Numbering is based upon the mature protein. **(D)** Octet BLI verification of U2S3 orthogonality when fused to the C-terminus of either the heavy or light chain of rituximab. Fc-wtNKG2D or Fc-iNKG2D.YA were captured with anti-human IgG Fc capture (AHC) biosensor tips then associated with a dilution series of MicAbody. The y-axes corresponding to binding responses were set to the same scale for all sensograms and Kd values for positive binding interactions are shown.

**Figure 2:**
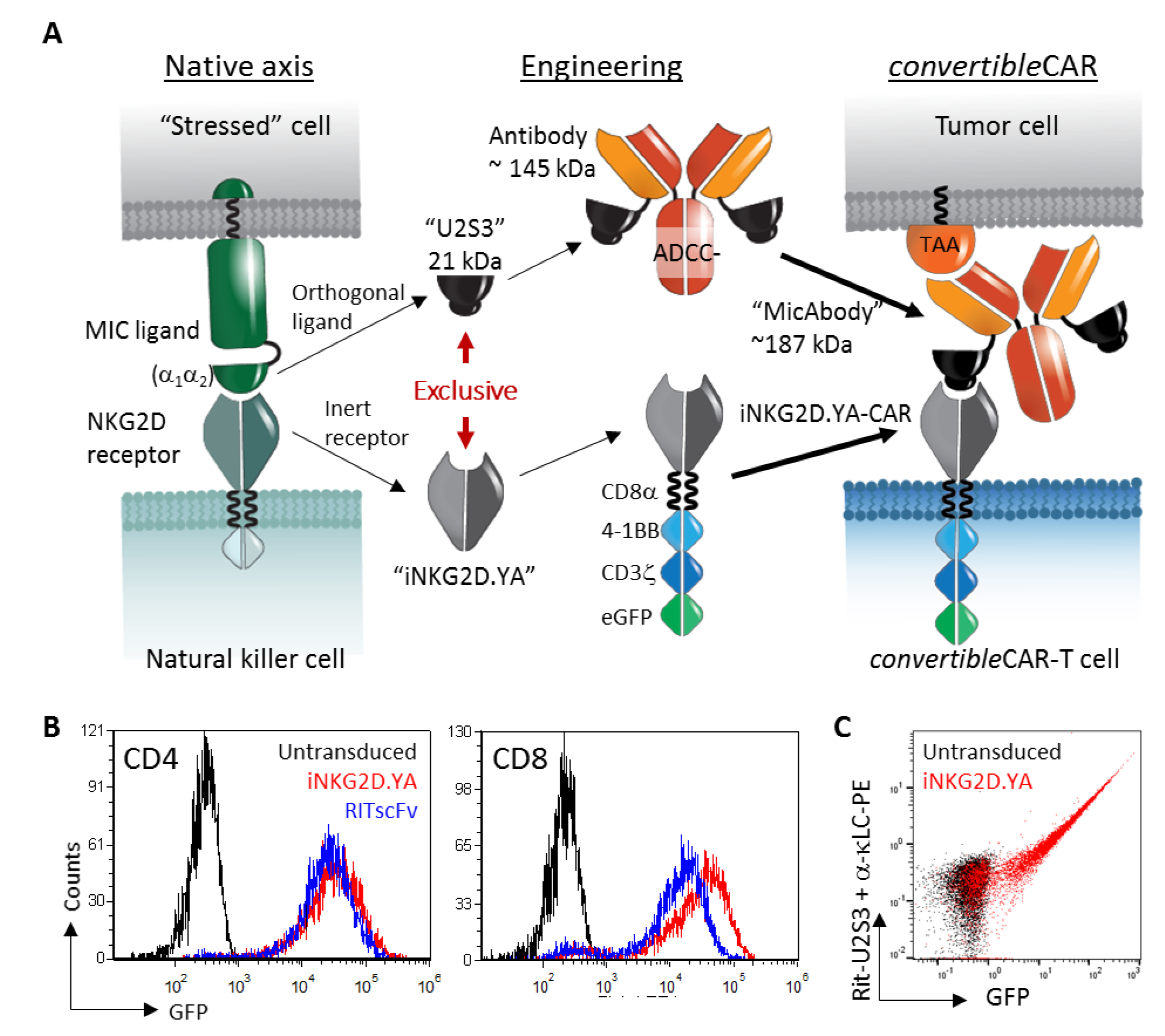
**(A)** Engineering schematic for the adaptation of the NKG2D-MIC axis into the *convertible*CAR components. iNKG2D.YA and U2S3 became modules of a second generation CAR receptor and bispecific adaptor molecule (MicAbody), respectively. The antigen-targeting MicAbody activates *convertible*CAR-expressing T cells only when the appropriate tumor-associated antigen (“TAA”) is expressed on a two-dimensional surface such as a tumor cell. **(B)** Representative example of high efficiency lentiviral transduction of the iNKG2D.YA-CAR into either CD4^+^ or CD8^+^ cells. Transduction efficiency varied between donors but >70% GFP^+^ yields were consistently achieved The RITscFv-CAR is shown for comparison and has the same architecture as the iNKG2D-CAR except that an scFv based upon the VH/VL domains of rituximab was used instead of iNKG2D.YA. **(C)** Surface expression of iNKG2D.YA-CAR was determined in CD8^+^ T cells by incubating cells with Rituximab.LC-U2S3 MicAbody followed by PE-conjugated mouse-anti-human kappa chain antibody staining. Untransduced T cells (black) are shown for comparison.

**Figure 3:**
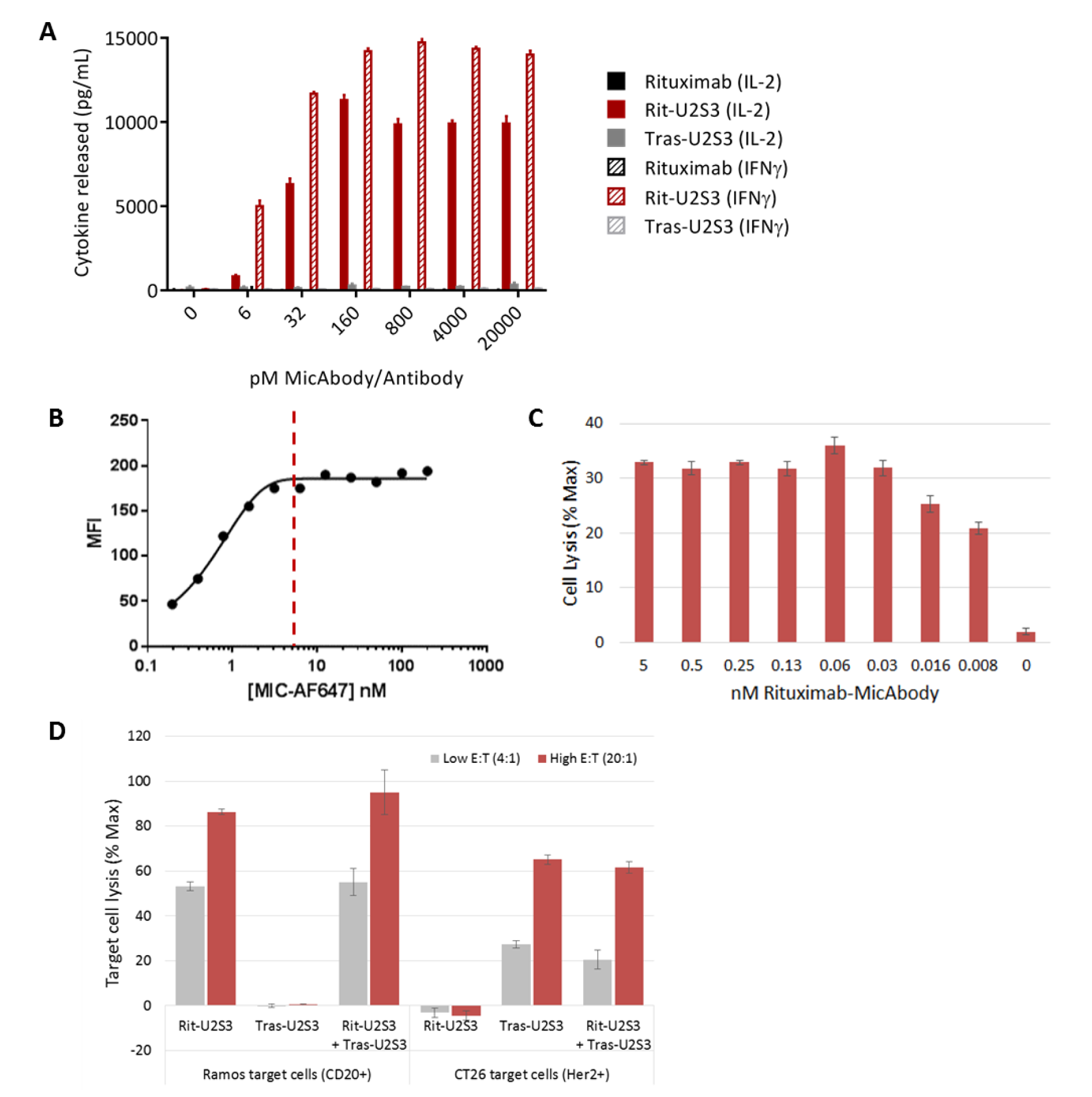
*In vitro* characterization of *convertible*CAR activity. **(A)** Ramos (CD20^+^) target cells were exposed to *convertible*CAR-CD8 cells at an E:T (effector to target ratio) of 5:1 and co-cultured with increasing concentrations of Rituximab antibody (ADCC-deficient), Rituximab.LC-U2S3 MicAbody, or Trastuzumab.LC-U2S3 MicAbody. After 24 hours, supernatants were harvested and IL-2 (solid bars) or IFNγ (hatched bars) quantified by ELISA. **(B)** *convertible*CAR-CD8^+^ cells were incubated with increasing concentrations of Alexa Fluor 647 conjugated Rituximab.LC-U2S3 for 30 minutes, the excess washed away, and the MFI quantified by flow cytometry. Dotted red line at 5 nM indicates inflection point at which receptors are maximally occupied. **(C)** *convertible*CAR-CD8 cells were armed with increasing concentrations of Rituximab.LC-U2S3 as described in (B) then co-incubated with calcein-loaded Ramos cells at an E:T of 10:1 for two hours after which the amount of released calcein was quantified. **(D)** iNKG2D.YA-CD8^+^ cells were pre-armed with 5 nM Rituximab.LC-U2S3, 5 nM Trastuzumab.LC-U2S3, or an equimolar mixture of 2.5 nM of each as described in (B) then exposed to calcein-loaded Ramos or CT26-Her2 cells at two indicated E:T ratios. The amount of calcein released was quantified after two hours. Error bars reflect standard deviation for replicates in the assay.

Candidate orthogonal variants were similarly identified for iNKG2D.AF and ELISAs comparing rituximab-LC fusions to Fc-wtNKG2D, Fc-iNKG2D.YA, and Fc-iNKG2D.AF identified four variants that selectively bound only iNKG2D.AF (SFig. 3B) with the U2R variant being the most selective. ELISAs comparing binding of Rituximab.LC-U2S3 and Rituximab.LC-U2R to both iNKG2D.YA and iNKG2D.AF confirmed that these two independently selected orthogonal ligands exclusively engaged the inert NKG2D variant for which it was evolved (SFig. 3C).

### Expression of iNKG2D.YA as a chimeric antigen receptor on T cells

Lentiviral transduction of iNKG2D.YA fused to 4-1BB, CD3ζ, and eGFP into primary human T cells efficiently generated *convertible*CAR-T cells with robust transgene expression on par with a rituximab-scFv based CAR construct (RITscFv-CAR) with the same hinge, transmembrane, and intracellular architecture (Fig. 2B). Surface staining of iNKG2D.YA with the Rituximab.LC-U2S3 MicAbody correlated strongly with GFP expression, suggesting a direct relationship between efficiency of CAR expression and presentation of iNKG2D on the T cell surface (Fig. 2C) and enabled quantification of a median of 21,000 molecules of iNKG2D.YA-receptors expressed on the surface (data not shown). Direct engagement of iNKG2D.YA-CAR receptors by incubation of *convertible*CAR-CD8^+^ cells to microtiter plates coated with wild-type or U2S3 ligands resulted in activation and liberation of IL-2 and IFNγ only with U2S3 while wtNKG2D-CAR bearing cells responded only to wild-type ligands confirming the selectivity of the orthogonal interaction in the context of T cells (SFig. 4A). Furthermore, activation of *convertible*CAR-T cell function was dependent upon the presence of the appropriate cognate ULBP2 variant. The iNKG2D.YA expressing or iNKG2D.AF expressing T cells only lysed Ramos (CD20^+^) target cells when armed with a MicAbody bearing its respective orthogonal ligand, i.e. U2S3 or U2R (SFig. 3D). Co-culture of Ramos cells alone was not sufficient to drive activation of *convertible*CAR-CD8^+^ cells. Instead the appropriate antigen-targeting MicAbody was required since neither rituximab antibody nor Trastuzumab.LC-U2S3 activated CAR cells whereas Rituximab.LC-U2S3 triggered maximum cytokine release in the 32-160 pM range. Additionally, cytokine release by *convertible*CAR-T cells plus Rituximab.LC-U2S3 MicAbody exceeded that of the RITscFv-CAR cells (SFig. 4B). These data demonstrated that the formation of an immunological synapse with CAR receptor clustering was required for robust activation of *convertible*CAR-T cells (SFig. 2A).

Staining of *convertible*CAR-CD8^+^ cells with a fluorescently labeled Rituximab.LC-U2S3 MicAbody revealed saturation of iNKG2D.YA-CAR receptors at 5 nM (Fig. 3B). In a co-culture experiment where *convertible*CAR-CD8^+^ cells were armed with decreasing amounts of Rituximab.LC-U2S3 prior to introduction to Ramos cells, target cell lysis remained close to maximum until the concentration used in arming was below 30 pM (Fig. 3C). There is therefore a two-order of magnitude difference between the concentrations of MicAbody required for full activation of function and complete occupancy of the receptors, suggesting a potential for arming the *convertible*CAR-T cell with more than one MicAbody to guide activity against multiple targets simultaneously. To directly test this, *convertible*CAR-CD8^+^ cells were armed with Rituximab.LC-U2S3, Trastuzumb.LC-U2S3 (targeting Her2), or an equimolar mixture of the two MicAbodies and exposed to either Ramos cells or CT26-Her2. Although CAR cells armed with a single MicAbody directed lysis to only tumor cells expressing the cognate antigen, dual-armed CARs targeted both tumor cell lines without any compromise in lytic potency (Fig. 3D).

### *convertible*CAR-T cells inhibit expansion of a disseminated B-cell lymphoma

The pharmacokinetics of both the HC and LC Rituximab-U2S3 MicAbodies (SFig. 5A) in NSG mice revealed a beta-phase that paralleled the parental antibody with the steeper alpha-phase of the MicAbodies attributed to retention of U2S3 binding to endogenous mouse wild-type NKG2D (data not shown). The LC-U2S3 fusion had a slightly longer terminal half-life than the HC-fused MicAbody, out-performed the HC fusion in an *in vitro* killing assay with Ramos target cells (SFig. 5B), and appeared to be more efficacious at early time points in suppressing Raji B cell lymphoma expansion in NSG mice (SFig. 5C, 5D). Accordingly, Rituximab.LC-U2S3 (Ritux-S3) was deployed in further experiments exploring dosing parameters for lymphoma control (SFig. 6). An intermediate Ritux-S3 dose of 20 μg was shown to be the most efficacious as it likely achieves a balanced density on both target and CAR cells for efficient generation of immunological synapses. Additionally, a higher frequency of Ritux-S3 administration of every two days versus every four days paired with a higher dose (10×10^6^) of *convertible*CAR-T cells resulted in the greatest suppression of tumor growth. Ritux-S3 alone was ineffective at tumor control while a graft-vs-tumor effect was consistently observed in both untransduced and *convertible*CAR only cohorts (SFigs. 6A, 6B; Fig. 4B). Ritux-S3 was detectable in the serum of mice throughout the course of the study with peak levels appearing earlier with more frequent dosing (SFig. 6B).

**Figure 4:**
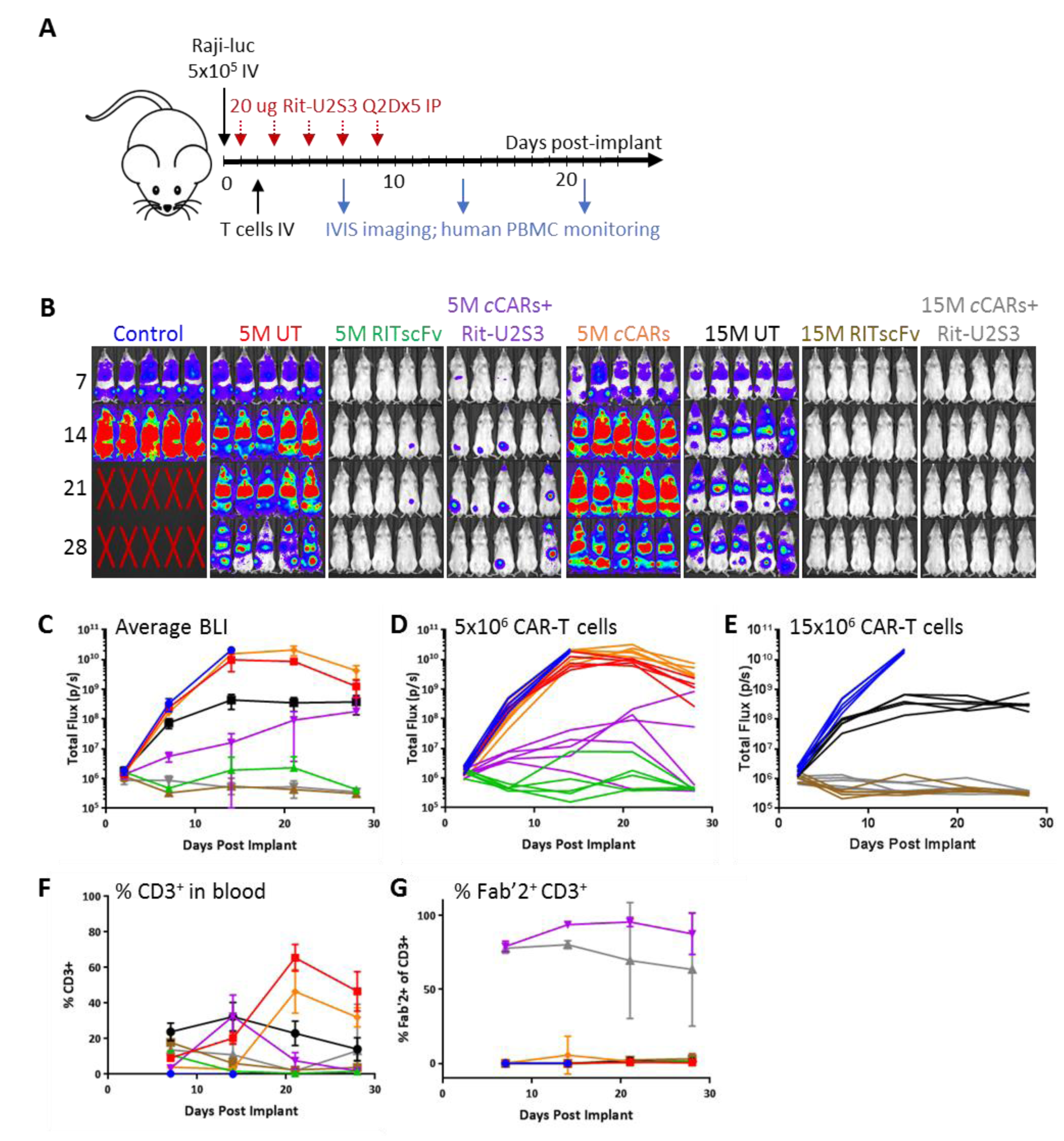
Control of a disseminated Raji B cell lymphoma in NSG mice. **(A)** Study design in NSG mice with Raji-luciferase cells implanted IV followed by treatment and monitoring as indicated. **(B)** IVIS imaging was performed 7, 14, 21, and 28 days post-implantation and all adjusted to the same scale. **(C)** Average luminescent output ±SD for each cohort along with individual animal traces for the groups that received **(D)** 5×10^6^ or **(E)** 15×10^6^ total T cells. T cell dynamics over the course of the study examining **(F)** human CD3^+^ cells in the blood and **(G)** bound MicAbody detected by anti-Fab’2. Shown are cohort averages ±SD. Color coding in all graphs matches the labeling on the luminescence images.

A Raji disseminated lymphoma model with optimized *convertible*CAR-T dosing was performed with 20 μg Ritux-S3 dosing every two days comparing 5×10^6^ (5M) to 15×10^6^ (15M) *convertible*CAR-T cells (Fig. 4A). As a positive control, RITscFv-CAR cells were also included and had been shown to have comparable *in vitro* Ramos killing potency as *convertible*CAR-T cells (data not shown and SFig. 4B). At 5M total T cells, both RITscFv-CAR and *convertible*CARs+Ritux-S3 were effective at controlling tumor (Fig. 4B). Although the average tumor bioluminescence signal was lower for the RITscFv-CAR cohort (Fig. 4C), and four of five mice in that cohort appeared to have cleared tumor tissue, three of five mice in the *convertible*CAR+Ritux-S3 cohort appeared cleared (Fig. 4D). When total infused CAR-T cell doses were increased to 15M cells, both RITscFv-CAR and *convertible*CAR+Ritux-S3 were able to completely block tumor expansion (Fig. 4B, 4C, 4D). In all studies, peak levels of peripheral human CD3^+^ T cells consistently appeared around seven days post-infusion with both scFv-CAR and *convertible*CAR-T cells having contracted in the majority of mice by 14 days (Fig. 4F). There was a delayed expansion of CD3^+^ cells in the untransduced and *convertible*CAR-only cohorts that was contemporary with the onset of the graft-versus-tumor response and likely the consequence of expansion of specific reactive clones. MicAbody associated *convertible*CAR-T cells were observed in the blood of mice in *convertible*CAR+Ritux-S3 cohorts (Fig. 4G) and these cells remain armed as long as sufficient plasma MicAbody is present (data not shown).

### *convertible*CAR-T cells inhibit growth of subcutaneously implanted Raji B-cells

Raji B-cells were implanted subcutaneously to assess the ability of the *convertible*CAR system to suppress growth of a solid tumor mass. Once tumors were established at 10 days, either 7×10^6^ (7M) or 35×10^6^ (35M) *convertible*CAR-Ts were administered after a single IV dose of 60 µg Ritux-S3 (Fig. 5A). Additionally, one cohort received 35M cells that were pre-armed with a saturating concentration of Ritux-S3 prior to administration but no additional MicAbody introduced injections. Administration of 7M *convertible*CAR-T cells along with Ritux-S3 (7M+MicAbody) resulted in reduced tumor size relative to *convertible*CAR-T cells alone (Figs. 5B, 5C). Furthermore, in the 35M+MicAbody cohort, tumor growth was completely suppressed. Tumor growth in the cohort that received 35M pre-armed cells was also inhibited. By two days post-infusion, *convertible*CAR-T cells that had been pre-armed did not have detectable surface associated MicAbody (Fig. 5E), a result of disarming likely due to a combination of activation-induced cell proliferation and the turnover armed receptors. Serum levels of Ritux-S3 were comparable at both CAR-T cell doses across the study and persisted through day 21 when it was detected at approximately 600 ng/mL (3.2 nM) (Fig. 5D) corresponding to high levels of armed peripheral CAR cells (Fig. 5E). By day 45 of the study, the cohort receiving 35M+MicAbody maintained relatively high CD3^+^ T cell numbers but were not well-armed with MicAbody while the 7M+MicAbody cohort did have cells that maintained surface-associated MicAbody. This suggested that as MicAbody levels fell below detectable limits in the plasma, CAR arming could not be maintained at high CAR-T cell levels. An alternative possibility is that the higher CD3^+^ cell numbers in the 35M+MicAbody cohort reflect expansion of a graft-vs-host subset of cells that do not express the CAR construct. However, the elevated CD3^+^ cell numbers were not seen in the 35M pre-armed cohort suggesting that this is not the case. In summary, pre-armed *convertible*CAR-Ts were able to exert a potent anti-tumor response that inhibited tumor expansion. Furthermore, *convertible*CAR-Ts were able to effectively control a solid lymphoma when an adequate *in vivo* level of *convertible*CAR-T cell arming was maintained.

**Figure 5:**
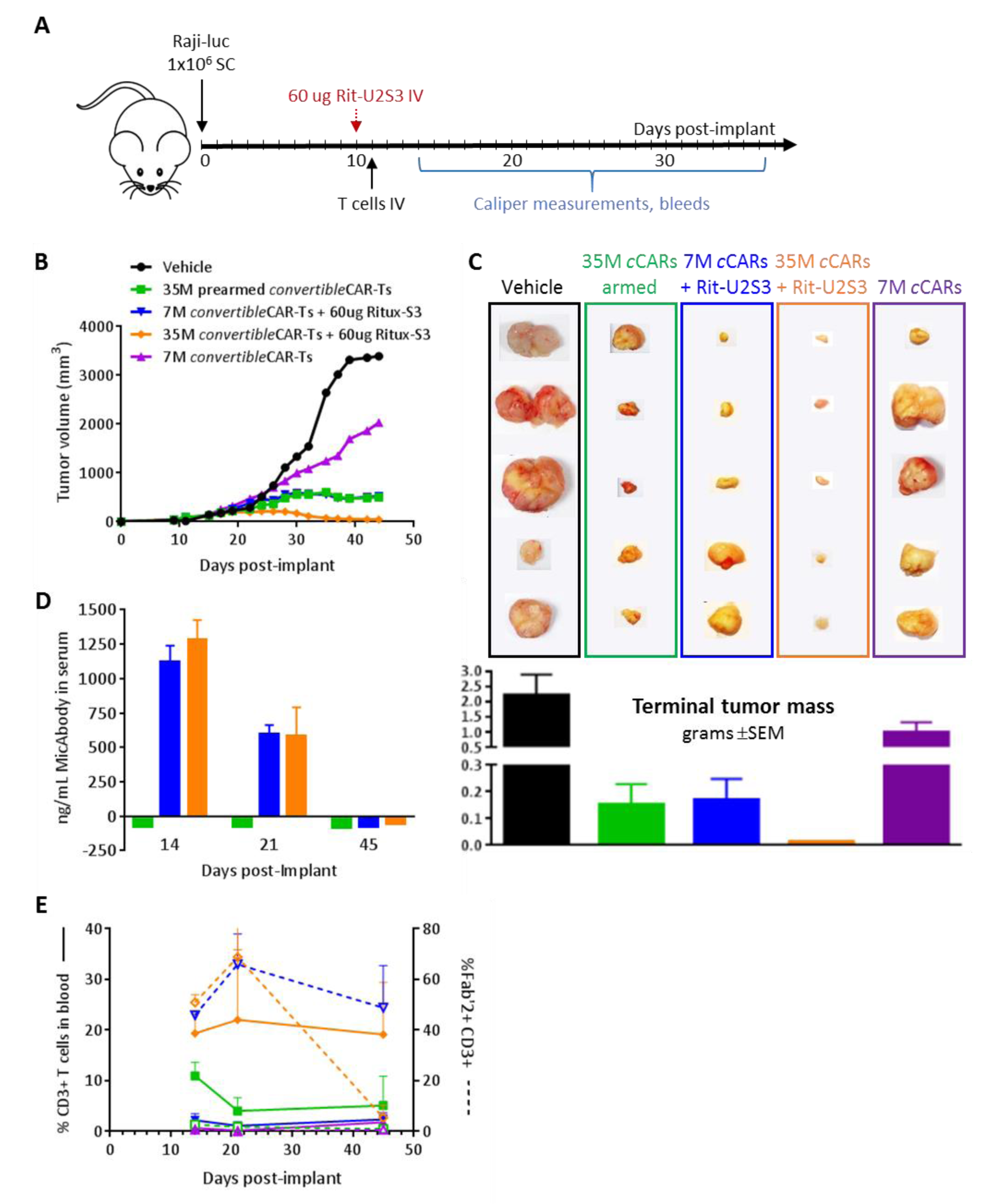
Control of subcutaneously implanted Raji tumors in NSG mice by *convertible*CAR-T cells. **(A)** Schematic of study design with treatment initiated at day 10 with Rituximab-MicAbody administration followed by CAR-T infusion the next day. **(B)** Average tumor volumes for each cohort. Tumors varied greatly in size within each group so error bars were not graphed. Two cohorts, blue and green lines, overlap and cannot be graphically distinguished beyond day 26. **(C)** Excised tumors at day 45 post-implantation with average tumor masses plotted in grams for each cohort. **(D)** Serum Rituximab-MicAbody levels. Error bars indicated ±SD. **(E)** CD3^+^ T cell dynamics in the blood and quantitation of the percentage of T cells with surface-associated MicAbody (Fab’2 staining). Groups are color coded to match across the different graphics.

### Selective delivery of modulatory reagents to *convertible*CAR-T cells

The privileged interaction between iNKG2D variants and their orthogonal ligands enables the selective delivery of agents to iNKG2D-CAR expressing cells simply by fusing them as payloads to the orthogonal ligands themselves. To demonstrate the utility of this feature two disparate applications were explored: targeted ablation utilizing the complement system and selective delivery of activating cytokines. In the first application, the U2R variant was fused to either the N- or C-terminus of the wild-type human IgG1 Fc-domain or to mutant Fc domains previously described as enhancing C1q binding - S267E/H268F/S324T/G236A/I332E (“EFTAE”)^18^ and K326A/E333A (“AA”)^19^ (SFig. 7A). Using just the Fc-portion, as opposed to a complete therapeutic antibody targeting an epitope-tagged CAR cell, avoids collateral effects from opsonization of non-iNKG2D expressing cells^20 21 22^. The enhanced C1q binding was confirmed by ELISA with relative order of Kd’s as EFTAE<AA<wt (SFig. 7B). While iNKG2D.AF-CAR cells were susceptible to killing by human complement in a manner that was dependent upon both concentration and C1q affinity (SFig. 7C), untransduced cells were unaffected. Interestingly, orientation of the U2R fusion was critical - N-terminal fusions, which orient the Fc in a manner consistent with an antibody were much more effective. Similar results were obtained with the U2S3 and iNKG2D.YA pairing (data not shown).

The potential ability of orthogonal ligands to deliver cytokines selectively to iNKG2D-CAR expressing cells has significant advantages to not only promote their expansion but also potentially leverage differential cytokine signaling to control T cell phenotype and function. As a general design principle, mutant cytokines with reduced binding to their natural receptor complexes were employed to reduce their engagement with immune cells not expressing the CAR and to minimize toxicity associated with wild-type cytokines. Additionally, cytokine fusions were kept monovalent to eliminate avidity-enhanced binding and signaling. To this end, the R38A/F42K mutations in IL-2 (mutIL2)^23^ and the V49D mutation in IL-15 (mutIL15)^24^ dramatically reduce binding to each cytokine’s respective Rα subunit while maintaining IL-2Rβ/γ complex engagement. Initial experiments using the iNKG2D.YA orthogonal variant U2S2 fused to either mutIL2 or mutIL15 promoted proliferation of iNKG2D.YA-CAR expressing cells but not those expressing wtNKG2D-CAR (SFig. 8A). Both cell populations expanded in the presence of the ULBP2.R81W variant, which does not discriminate between wtNKG2D and iNKG2D.YA. Direct fusion to the ligand or via a heterodimeric Fc linkage^25^ (e.g. U2S2-hFc-mutIL2) promoted expansion of GFP^+^*convertible*CAR-T cells to densities above the untransduced cells present (SFig. 8B), and these expanded *convertible*CAR-T cells maintained their cytolyic capabilities (data not shown). Engagement of iNKG2D.YA by ligand either in the form of a MicAbody (Fig. 3A) or a monovalent U2S3-hFc (without a cytokine payload), by mutIL2 alone, or hFc-mutIL2 without U2S3 were all insufficient to drive proliferation of *convertible*CAR-CD8 cells (data not shown). Flow cytometry characterization of STAT3 and STAT5 phosphorylation (pSTAT3 and pSTAT5) revealed that exposure to wild-type IL-2 or IL-15 resulted in an increase of pSTAT3 and pSTAT5 in both untransduced as well as *convertible*CAR-CD8 cells (SFig. 8C). Treatment of untransduced cells with U2S3-hFc-mutIL2 resulted only in a minimal shift in pSTAT5 relative to the no cytokine control, consistent with mutIL2’s retention of IL-2Rβ/γc binding. The *convertible*CAR-CD8 cells responded to both U2S3-hFc-mutIL2 and U2S3-hFc-mutIL15 with an increase in pSTAT5 levels via γ-chain activation of JAK3. Unlike wild-type cytokines, no increase in pSTAT3 signal was observed, indicating a reduction in JAK1 activation through IL-2Rβ^26^ in both scenarios as a consequence of disruption of Rα binding, a hypothesis supported by IL-15Rα’s role in increasing the affinity of IL-15 for IL-2Rβ^27^. The kinetics of responses U2S3-hFc-mutIL2 and U2S3-hFc-mutIL15 were nearly identical, indicating functional redundancy in their mutant forms (SFig. 8D).

U2S3-hFc-mutIL2 was shown to have an *in vivo* PK half-life of a few days (SFig. 8E). *convertible*CAR-T cells administered to NSG mice in the absence of tumor underwent a homeostatic expansion, peaking at three days followed by contraction. Three injections of U2S3-hFc-mutIL2 staged one week apart resulted in a dramatic expansion of human T cells in the peripheral blood (Fig. 6A) and T cell numbers contracted after cessation of U2S3-hFc-mutIL2 support. While CD4^+^ T cells did expand somewhat (data not shown), the majority of expansion was by CD8^+^ T cells. In parallel with expansion, the proportion of GFP^+^ CD8^+^ T cells increased to 100% demonstrating selective expansion of iNKG2D-CAR expressing cells but not untransduced cells (Fig. 6B).

**Figure 6:**
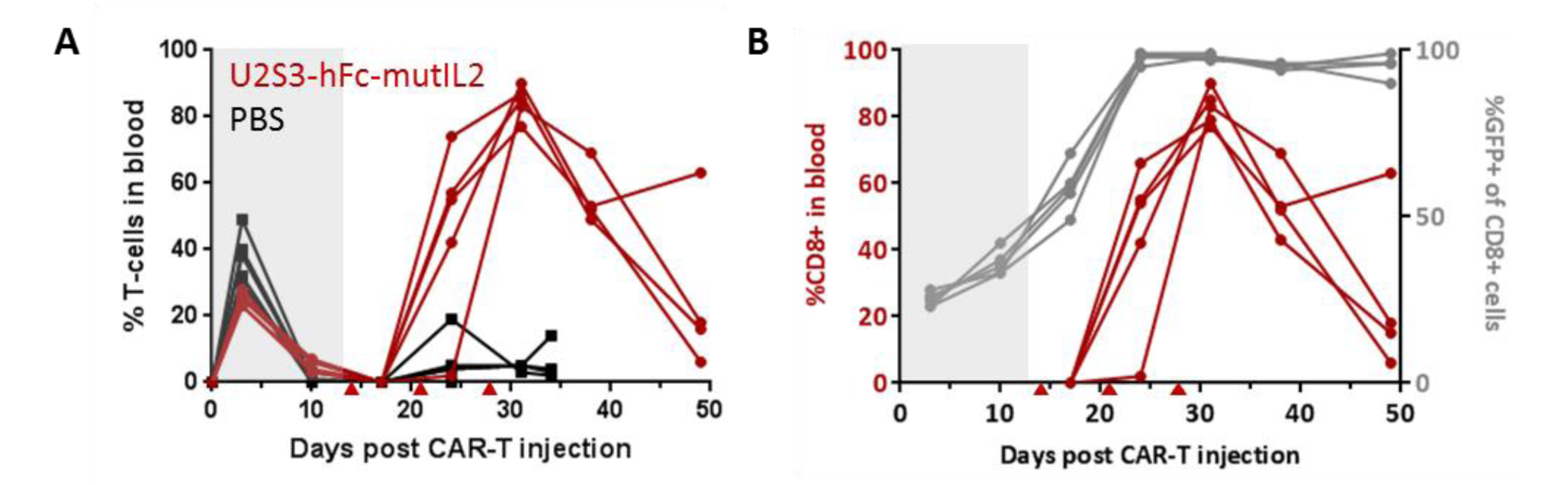
*In vivo* response of *convertible*CAR-T cells to U2S3-hFc-mutIL2. **(A)** NSG mice were injected with 7×10^6^ total iNKG2D-transduced cells (CD4:CD8 1:1). After contraction of T cells at 14 days, mice were injected with 30 μg U2S3-hFc-mutIL2 or PBS once per week – indicated by red triangles - and T cell dynamics monitored by flow. Shown is the % of human CD3^+^ T cells in the peripheral blood with each trace corresponding to an individual mouse. **(B)** Plots for the expansion of CD8^+^ cells as well as the increase in proportion of GFP^+^ (CAR-expressing) cells upon U2S3-mutIL2 treatment.

The effect of U2S3-hFc-mutIL2 on normal human PBMCs from three donors was explored *in vitro* by exposure to increasing concentrations of the agent for four days followed by flow-based quantification of cells positive for the proliferative marker Ki-67 (SFig. 9). In addition to the -mutIL2 fusion, a wild-type IL2 fusion (U2S3-hFc-wtIL2) was included to directly demonstrate that the reduction in mutIL2 bioactivity was a consequence of the mutations employed and not the fusion format itself.

The CD4^+^ and CD8^+^ T cells responded robustly to both anti-CD3 and wild-type IL-2 positive controls as well as to the lowest dose of U2S3-hFc-wtIL2. Proliferative responses to U2S3-hFc-mutIL2 occurred in a dose-dependent manner with expansion observed across donors at levels above 300 IUe/mL but not achieving levels comparable to those of the IL-2 positive control until 30,000 IUe/mL. Treg responses were comparable to those of CD4^+^ and CD8^+^ cells with the exception of cells from one donor (who additionally had a muted response to anti-CD3 stimulation) that responded to U2S3-hFc-mutIL2 at a lower concentration than the other donors. Taken together, these data support the hypothesis that normal human PBMCs do not respond to U2S3-hFc-mutIL2 except at super-physiologic levels which potentially provides a wide dosing window for selective delivery of ligand-fused mutIL2 to *convertible*CAR cells while minimizing toxicity and Treg activation.

## DISCUSSION

We have described the engineering of a privileged receptor-ligand (iNKG2D.YA and U2S3) pairing comprised of human components for a highly adaptable CAR, resulting in a versatile and broadly controllable platform. The iNKG2D.YA-CAR receptor itself is held invariant on T cells with CAR function readily directed to potentially any antigen of interest by virtue of attaching the orthogonal ligand to the appropriate antigen-recognizing antibody. In this manner, the same *convertible*CAR-T cells can be retargeted as needed if, for example, the original tumor antigen becomes downregulated during the course of therapy. This targeting flexibility is not limited to sequential engagement of antigens, but can also be multiplexed to simultaneously direct T cells to more than one antigen in order to reduce the likelihood of tumor escape by antigen loss, address the issue of heterogeneity of intratumoral antigen expression, or even simultaneously target tumor and suppressive cellular components of the tumor microenvironment. Typical scFv-CAR cells are generally committed to a fixed expression level of a receptor which reduces their ability to discriminate between antigen levels present on healthy versus aberrant cells. The use of MicAbodies provides an opportunity to differentially engage *convertible*CAR-T cells in order to achieve a therapeutic index that reduces the risk of severe adverse events.

The use of the privileged receptor-ligand interaction for delivery of payloads specifically to iNKG2D-bearing cells without additional cellular engineering is another significant advantage. The capability of harnessing interleukin functions to drive expansion and activation, prevent exhaustion, or even promote suppression in a controlled and targeted manner could have beneficial consequences for efficacy and safety. Introduction of cytokine-ligand fusions during CAR manufacturing could address qualitative and quantitative limitations of patient T cells and their administration post-CAR infusion could expand the number of CAR-T cells and their persistence which, with CD19-CAR therapies, is correlated positively with response rates^28^. Most CAR therapies require a preconditioning lymphodepletion regimen to promote engraftment and expansion of CAR cells, one rationale being that it provides a more verdant immunological setting for CARs to expand^29^. Robust and controllable *convertible*CAR-T expansion in patients may supplant the need for lymphodepletion, allowing for retention of endogenous immune functions that are fully competent to support the initial *convertible*CAR-mediated anti-tumor activity. Another clinical strategy might be to deliver cytokine-ligand fusions to bolster *convertible*CAR-T function, possibly with a cycling regimen to reduce T cell exhaustion and promote the maintenance of memory T cells^30^. And lastly, as CARs have been demonstrated to persist in humans for years post-infusion^31^, the ability to recall resident *convertible*CAR-Ts to attack primary or secondary malignancies (either with the original targeting MicAbody or a different one) without having to re-engineer or generate a new batch of CAR cells should be highly advantageous. Unlike scenarios where CARs have been engineered to constitutively express cytokines^32 33^, delivery of cytokines exclusively to *convertible*CAR-T cells can be modulated depending upon the manufacturing or clinical needs.

By design, each component of the *convertible*CAR system – the iNKG2D-based CAR receptor and the MicAbody (which is ADCC-deficient) – are functionally inert on their own. This has significant advantages during manufacturing, particularly in the context of indications such as T cell malignancies where traditional scFv-based CARs encounter expansion hurdles due to fratricide^34^. Additionally, it provides enhanced control of CAR function during treatment. We have demonstrated that *convertible*CAR-T cells can be armed with MicAbody prior to administration to provide an initial burst of anti-tumor activity on par with traditional scFv-CARs, an aspect that will be critical for first-in-human safety studies. In addition to activation-induced replication, these cells also internalize their engaged CAR receptors in a manner consistent with what has been observed with other 4-1BB/CD3zeta scFv-CARs^35^ (data not shown). As a consequence of these two processes, *convertible*CAR-T cells will rapidly disarm after initial expansion and target engagement, which then provides an opportunity rearm and re-engage in a manner controlled by MicAbody dosing.

In addition to the iNKG2D-U2S3 pairing based upon ULBP2, we have also identified high-affinity orthogonal MicA and ULBP3 variants to iNKG2D.YA that are non-redundant in their amino acid compositions through the helix 4 domain (data not shown). The fact that a convergence of sequence was not observed, despite the conserved tertiary structure of their MIC α_1_-α_2_ domains, indicates that there are many solutions for orthogonality. Additionally, we have engineered a completely independent iNKG2D.AF and U2R pairing suggesting that there is opportunity to explore a breadth of inert NKG2D variants that, in combination with mutational space afforded by the MIC ligands, can lead to the identification of additional orthogonal receptor-ligand pairings. Having mutually exclusive receptor-ligand pairs would enable, for example, their introduction into distinct cell populations (e.g. CD4 and CD8 T-cells) to differentially engage them as needed. Furthermore, within the same cell, the two iNKG2D variants could be expressed with split intracellular signaling domains to provide dual antigen-dependent activation to enhance on-tumor selectivity^36 37^. Alternatively, the two iNKG2D variants could be differentially linked to either activating or immunosuppressive domains to enhance the discriminatory power of the T cells between tumors or healthy tissue, respectively^38^.

In summary, the *convertible*CAR system has demonstrated capabilities to not only be readily targeted to different cell-surface antigens but can also be selectively engaged exogenously to drive cell expansion. The privileged receptor-ligand interaction that has been developed is agnostic to cell type and can, in principle, be engineered into any cell of interest as long as the cell-appropriate signaling domains are provided. Additionally, the adoptive cellular therapy field is aggressively pursuing the development of allogeneic cells to bring down the time, complexity, and cost of manufacturing to provide a more consistent, readily accessible product^39 28^. A highly adaptable CAR system would be powerfully synergistic with allogenic efforts and once a truly universal allogeneic CAR system has been validated, the therapeutic field then becomes characterized by the relative ease of developing and implementing a library of adaptor molecules from which personalized selections can be made. This strategy also broadens the potential areas of application to any pathogenic cell with a targetable surface antigen. For example, we have recently demonstrated the utility of *convertible*CAR platform in the *ex vivo* reduction of latent reservoirs in the blood of HIV patients using a cocktail of MicAbodies derived from anti-HIV broadly neutralizing antibodies^40^.

## MATERIALS AND METHODS

### Cloning, expression, and purification

The wild-type ectodomain of NKG2D (UniProtKB P26718, residues 78-216; https://www.uniprot.org) was expressed as a fusion to the C-terminus of human IgG1 Fc via a short factor Xa recognizable Ile-Glu-Gly-Arg linker (Fc-wtNKG2D). Inert NKG2D variants comprising either a single Y152A (iNKG2D.YA) or double Y152A/Y199A substitution (iNKG2D.AF)^16^ were generated by PCR-mediated mutagenesis or synthesized (gBlocks®, IDT). DNA constructs for Fc-NKG2D molecules were expressed in Expi293^TM^ cells (Thermo Fisher Scientific) and dimeric secreted protein was purified by Protein A affinity chromatography (Pierce^TM^ #20334, Thermo Fisher). Eluted material was characterized and further purified by size-exclusion chromatography (SEC) on an ÄKTA Pure system using Superdex 200 columns (GE Life Sciences). Correctly assembled, size-appropriate monomeric material was fractionated into phosphate-buffered saline (PBS).

The α_1_-α_2_ domains of human MICA*001 (UniProtKB Q29983, residues 24-205), MICB (UniProtKB Q29980.1, 24-205), ULBP1 (UniProtKB Q9BZM6, 29-212), ULBP2 (UniProtKB Q9BZM5, 29-212), ULBP3 (UniProtKB Q9BZM4, 30-212), ULBP5 (NCBI accession NP_001001788.2, 29-212), ULBP6 (UniProtKB, 29-212) were cloned with a C-terminal 6x-His tag. Monomeric protein was purified from Expi293^TM^ supernatants Ni-NTA resin (HisPur^TM^, Thermo Fisher) and eluted material exchanged into PBS with Sephadex G-25 in PD-10 Desalting Columns (GE Life Sciences).

MIC ligands and orthogonal variants were cloned by ligation-independent assembly (HiFi DNA Assembly Master Mix, NEB #E2621) as fusions to the C-terminus of either the kappa light-chain or the heavy-chain of human IgG1 antibodies via either an APTSSSGGGGS or GGGS linker, respectively. Additionally, D265A/N297A (Kabat numbering) mutations were introduced into the CH2 domain of the heavy chain of all antibody and MicAbody clones to eliminate antibody-dependent cell cytotoxicity (ADCC) function^41^. Heavy- and light-chain plasmid DNAs (in the mammalian expression vector pD2610-V12 (ATUM) for a given antibody clone were co-transfected into Expi293^TM^ cells and purified by Protein A. For any monoclonal antibody fusion generated, the appropriate VL or VH domains were swapped into either the kappa light-chain or ADCC-deficient IgG1 heavy-chain.

### Engineering of inert NKG2D, identification of orthogonal ligand variants, and MicAbody characterization

Bio-layer interferometry (BLI) with the FortéBio Octet system (Pall FortéBio LLC) was implemented to validate loss of wild-type MIC ligand binding by iNKG2D. Fc-wtNKG2D, Fc-iNKG2D.YA, or Fc-iNKG2D.AF was captured on anti-human IgG Fc capture (AHC) biosensor tips and association/dissociation kinetics monitored in a titration series of monomeric MIC-His ligands. Additionally, ELISA (enzyme-linked immunosorbent assay) binding assays were performed with MICA-Fc, MICB-Fc, ULBP1-Fc, ULBP2-Fc, ULBP3-Fc, or ULBP4-Fc (R&D Systems) coated onto microtiter plates, a titration of biotinylated Fc-wtNKG2D or Fc-iNKG2D.YA, detected with streptavidin-HRP (R&D Systems #DY998), and developed with 1-Step Ultra TMB ELISA (Thermo Fisher #34208).

Phage display was employed to identify orthogonal ULBP2 α_1_-α_2_ variants that exhibited exclusive binding to either iNKG2D.YA or iNKG2D.AF. Synthetic NNK DNA libraries were generated targeting the codons of helix 2 (residues 74-78, numbering based upon mature protein) or helix 4 (residues 156-160) that in the bound state are positioned in close proximity to the Y152 positions on the natural NKG2D receptor^42^. Libraries exploring helix 2 alone, helix 4 alone, or the combination were cloned as fusions to the pIII minor coat protein of M13 phage, and phage particles displaying the mutagenized α_1_-α_2_ domain variants were produced in SS320 *E.coli* cells according to standard methods^43 44^. These α_1_-α_2_ phage phage libraries were captured with either biotinylated Fc-iNKG2D.YA or Fc-iNKG2D.AF protein (EZ-Link^TM^ NHS-Biotin Kit, Thermo Fisher #20217) and enriched by cycling through four rounds of selection with increasing concentrations of non-biotinylated Fc-wtNKG2D competitor. Positive phage clones were verified for preferential binding to plate-bound Fc-iNKG2D.YA or Fc-iNKG2D.AF versus Fc-wtNKG2D by spot ELISA and bound phage detected with biotinylated M13 phage coat protein monoclonal antibody E1 (Thermo Fisher # MA1-34468) followed by incubation with streptavidin-HRP.

Phage variants were sequenced then cloned as human IgG1 monoclonal antibody fusions (see above) for additional validation. To confirm that selectivity of orthogonal variants was maintained in the bivalent MicAbody format, ELISAs wells were coated with 1 μg/mL Fc-wtNKG2D, Fc-iNKG2D.YA, or Fc-iNKG2D.AF and bound MicAbody detected with an HRP-conjugated mouse-anti-human kappa chain antibody (Abcam #ab79115). Affinity of both monomeric and antibody-fused ULBP2 variants was also determined by Octet analysis as described above.

### CAR construction, lentiviral production, human primary T cell isolation, and lentiviral transduction

Human-codon optimized DNA (GeneArt, Thermo Fisher) comprising the CD8α-chain signal sequence, NKG2D variant, CD8α hinge and transmembrane domains, 4-1BB, CD3ζ, and eGFP were cloned into the pHR-PGK transfer plasmid for second generation Pantropic VSV-G pseudotyped lentivirus production along with packaging plasmids pCMVdR8.91 and pMD2.G^45^. The VH and VL domains of rituximab separated by a (GGGGS)3 linker were substituted for the NKG2D module to generate the rituximab scFv-based CAR (RITscFv-CAR). For each batch of lentivirus produced, 6×10^6^ Lenti-X 293T (Takara Bio #632180) cells were seeded in a 10 cm dish the day prior to transfection. Then 12.9 µg pCMVdR8.91, 2.5 µg pMD2.G and 7.2 µg of the pHR-PGK-CAR constructs were combined in 720 µl Opti-MEM^TM^ (Thermo Fisher #31985062) then mixed with 67.5 µl of Fugene HD (Promega Corp. E2311), briefly vortexed, and incubated at room temperature for 10 minutes before adding to the dish of cells. After two days, supernatants were collected by centrifugation and passed through 0.22 µm filters. 5X concentrated PEG-6000 and NaCl were added to achieve final concentrations of 8.5% PEG-6000 (Hampton Research #HR2-533) and 0.3 M NaCl, incubated on ice for two hours, then centrifuged at 4⁰C for 20 minutes. Concentrated viral particles were resuspended in 0.01 volume of PBS, and stored frozen at −80⁰C.

For primary human T cell isolation, a Human Peripheral Blood Leuko Pak (Stemcell Technologies #70500.1) from an anonymous donor was diluted with an equivalent volume of PBS + 2% FBS, then centrifuged at 500 x g for 10 minutes at room temperature. Cells were resuspended at 5×10^7^ cells/ml in PBS + 2% FBS and CD4^+^ or CD8^+^ cells enriched by negative selection (Stemcell EasySep^TM^ Human CD4 T Cell Isolation Kit #17952 or EasySep Human CD8 T Cell Isolation Kit #17953) by addition of 50 µl of isolation cocktail per ml of cells and incubating for five minutes at room temperature. Subsequently, 50 µl of RapidSpheres^TM^ were added per ml of cells and samples topped off (to each 21 mL cells, 14 mL of PBS). Cells were isolated for 10 minutes with an EasySEP^TM^ magnet followed by removal of buffer while maintaining the magnetic field. Enriched cells were transferred into new tubes with fresh buffer and the magnet reapplied for a second round of enrichment after which cells were resuspended, counted, and cryopreserved at 10-15×10^6^ cells/cryovial (RPMI-1640, Corning #15-040-CV; 20% human AB serum, Valley Biomedical #HP1022; 10% DMSO, Alfa Aesar #42780).

To generate CAR-T cells, one vial of cryopreserved cells was thawed and added to 10 ml T cell medium “TCM” (TexMACS medium, Miltenyi 130-097-196; 5% human AB serum, Valley Biomedical #HP1022; 10 mM neutralized N-acetyl-L-Cysteine; 1X 2-mercaptoethanol, Thermo Fisher #21985023, 1000X; 45 IUe/ml human IL-2 IS “rhIL-2”, Miltenyi #130-097-746) added at time of addition to cells. Cells were centrifuged at 400 x g for 5 minutes then resuspended in 10 ml TCM and adjusted to 1×10^6^/ml and plated at 1 ml/well in a 24 well plate. After an overnight rest 20 μL of Dynabeads™ Human T-Activator CD3/CD28 (Thermo Fisher #1131D) were added per well and incubated for 24 hours. Concentrated lentiviral particles (50 µL) were added per well, cells incubated overnight, then transferred to T25 flasks with an added 6 ml TCM. After three days of expansion, Dynabeads were removed (MagCellect magnet, R&D Systems MAG997), transduction efficiency assessed by flow cytometry for GFP, back-diluted to 5×10^5^ cells/mL, and cell density monitored daily to ensure they did not exceed 4×10^6^ cells/ml. When necessary, surface expression of iNKG2D was correlated with GFP expression using a MicAbody and detecting with PE-anti-human kappa chain (Abcam #ab79113) or by directly conjugating the Rituximab-MicAbody to Alexa Fluor 647 (Alexa Fluor Protein Labeling Kit #A20173, Thermo Fisher). The amount of iNKG2D expression on the surface of *convertible*CAR-CD8 cells was quantified using Alexa Fluor 647 conjugated Rituximab-MicAbody, and median fluorescence intensity was correlated with Quantum^TM^ MESF 647 beads (Bangs Laboratories #647). All flow cytometry was performed on either Bio-Rad S3e Cell Sorter or Miltenyi MACSQuant Analyzer 10 instruments.

### Cell lines and *in vitro* assays

Ramos human B cell lymphoma cells (ATCC #CRL-1596) were cultured in RPMI supplemented with 20 mM HEPES and 10% FBS. The mouse colon carcinoma line CT26 transfected to express human Her2 were a kind gift from Professor Sherie Morrison (UCLA).

For calcein-release assays, tumor cells were centrifuged and resuspend in 4 mM probenecid (MP Biomedicals #156370) + 25 µM calcein-AM (Thermo Fisher #C1430) in T cell medium at 1-2×10^6^ cells/ml for one hour at 37⁰C, washed once, and adjusted to 8×10^5^ cells/ml. CD8^+^ CAR-T cells were pelleted and resuspended in 4 mM probenecid with 60 IUe/ml IL-2 in TCM at 4×10^6^ cells/mL then adjusted according to the desired effector:target ratio (unadjusted for transduction efficiency). 25 µL target cells were plated followed by 25 µL medium or diluted MicAbody. Then 100 μL medium (minimum lysis), medium + 3% Triton-X 100 (maximum lysis), or CAR-T cells were added and plates incubated at 37⁰C for two hours. Cells were pelleted and 75 μL supernatant transferred to black clear-bottom plates and fluorescence (excitation 485 nm, emission cutoff 495 nm, emission 530 nm, 6 flashes per read) acquired on a Spectramax M2e plate reader (Molecular Devices). For experiments with armed *convertible*CAR-CD8^+^s, T cells were pre-incubated at 37⁰C with either saturating (5 nM) or a titration of MicAbody for 30 minutes before washing to remove unbound MicAbody and co-culturing with calcein-loaded target cells.

In order to quantify the target-dependent activation of T-cells, experiments were set up as described above except that calcein-preloading was omitted and assays set up in T cell medium without IL-2 supplementation. After 24 hours co-culture, supernatants were harvested and stored at −80°C until the amount of liberated cytokine could be quantified by ELISA MAX^TM^ Human IL-2 or Human IFN-g detection kits (BioLegend #431801 and #430101).

To generate a MicAbody binding curve to iNKG2D.YA-CAR expressing T-cells, Rituximab.LC-U2S3 was labeled with Alexa Fluor 647. 3×10^5^ *convertible*CAR-CD8^+^ cells were plated in 96-wells V-bottom plates and incubated with labeled MicAbody for 30 minutes at room temperature in a final volume of 100 μL RPMI + 1% FBS with a titration curve starting at 200 nM. Cells were then rinsed and median fluorescence intensity determined for each titration point by flow cytometry.

### Animal studies

All animal studies were conducted by ProMab Biotechnologies, Inc. (Richmond, CA) with the exception of the U2S3-hFc-mutIL2 pharmacokinetic (PK) sampling which was conducted by Murigenics (Vallejo, CA). For PK analysis of serum levels of MicAbodies, six-week old female NSG mice (NOD.Cg-*Prkdc^scid^ IL2rg^tm1Wjl^*/SzJ, The Jackson Laboratory # 005557) were injected intravenously (IV) with 100 μg of either parent rituximab antibody (ADCC-defective), heavy-chain U2S3 fusion of rituximab (Rituximab.HC-U2S3), or light-chain fusion (Rituximab.LC-U2S3). Collected sera were subjected to ELISA by capturing with human anti-rituximab idiotype antibody (HCA186, Bio-Rad Laboratories), detected with rat-anti-rituximab-HRP antibody (MCA2260P, Bio-Rad), and serum levels interpolated using either a rituximab or Ritxumab-U2S3 standard curve. PK analysis of U2S3-hFc-mutIL2 was performed in NSG mice by IP injection of 60 μg followed by regular serum collection. Samples were examined by ELISA capturing with Fc-iNKG2D and detecting with biotinylated rabbit-anti-human IL-2 polyclonal antibody (Peprotech #500-P22BT) followed by incubation with streptavidin-HRP. Half-lives were calculated in GraphPad Prism based upon the beta-phase of the curve using a nonlinear regression analysis, exponential, one-phase decay analysis with the plateau constrained to zero.

For disseminated Raji B cell lymphoma studies, six-week old female NSG mice were implanted IV with Raji cells stably transfected to constitutively express luciferase from *Luciola italica* (Perkin Elmer RediFect Red-FLuc-GFP # CLS960003). Initiation of treatment administration is detailed in each *in vivo* study figure. For all experiments, CD4 and CD8 primary human T cells were independently transduced, combined post-expansion at a 1:1 mixture of CD4:CD8 cells without normalizing for transfection efficiency between cell types or CAR constructs, and the mixture validated by flow cytometry prior to IV injection. Administration of MicAbody or control antibody was by the intraperitoneal (IP) route unless otherwise specified, and *in vivo* imaging for bioluminescence was performed with a Xenogen IVIS system (Perkin Elmer). Animals were bled regularly to monitor human T cell dynamics by flow cytometry, staining with APC Anti-Human CD3 (clone OKT3, 20-0037-T100, Tonbo Biosciences), monitoring GFP, and examining cell-associated MicAbody levels with biotinylated Anti-Human F(ab’)2 (109-066-097, Jackson ImmunoResearch Laboratories Inc.). Serum ELISAs to monitor MicAbody levels was performed as described above.

Subcutaneous Raji B cell tumor studies were performed in NSG mice implanted with 1×10^6^ Raji cells in matrigel on the right flank and therapy initiated when tumors reached 70-100 mm^3^. For the cohort that received armed *convertible*CAR-T cells, the cells were incubated with 5 nM Rituximab.LC-U2S3 MicAbody *ex vivo* for 30 minutes at room temperature before washing and final mixing to achieve the desired 1:1 CD4:CD8 ratio and cell concentration. Arming was confirmed by flow cytometry with the biotinylated Anti-Human F(ab’)2 antibody and revealed a strong correlation between GFP and F(ab’)2 MFIs. These mice did not receive a separate MicAbody administration. Caliper measurements were regularly taken to estimate tumor volume, and terminal tumor masses were weighed.

### Targeted delivery of complement C1q to cells expressing iNKG2D.AF-CAR

To generate Fc reagents with enhanced complement binding and targeted delivery to the T cells expressing iNKG2D.AF, the orthogonal ligand was cloned as a fusion to either the N-(U2R-Fc) or C-terminus (Fc-U2R) of human IgG1 Fc via a GGGS linker with the Fc including the hinge, CH2, and CH3 domains. In addition to the wild-type Fc, the K326A/E333A^19^ (Kabat numbering, “AA”) and S267E/H268F/S324T/G236A/I332E^18^ (“EFTAE”) C1q-enhanced binding mutation sets were explored. All were expressed in Expi293T cells, purified, and fractionated as described above. Confirmatory ELISAs were performed by capturing with Fc-NKG2D.AF followed by binding U2R/Fc-variant fusions at 1 μg/mL concentration, titrating in human-C1q protein (Abcam #ab96363), then detecting with polyclonal sheep-anti-C1q-HRP antibody (Abcam #ab46191). Complement-dependent cytotoxicity (CDC) assays were performed by iQ Biosciences (Berkeley, CA). Briefly, 5×10^4^ CD8^+^ cells from an NKG2D.AF-CAR transduction were plated in 96-well plates and incubated for three hours with a serial dilution of each U2R/Fc-variant fusion, in triplicate, in the presence of normal human serum complement (Quidel Corporation) at a final concentration of 10% (v/v). Cells were then harvested and resuspended with SYTOX^TM^ Red dead cell stain (Thermo Fisher) at a final concentration of 5 μg/mL and analyzed by flow cytometry. EC50 values for cytotoxicity were calculated in GraphPad prism fitted to a non-linear regression curve.

### Targeted delivery of mutant-IL2 to T cells expressing iNKG2D-CAR

To generate a reagent that was monomeric for the U2S3 ligand, monomeric for a mutant IL-2 with significantly reduced IL-2Rα binding (mutIL2, R38A/F42K)^23 46^ yet retained serum stability, a heterodimeric Fc strategy was employed^25^. U2S3 was fused to the N-terminus of the Fc-hinge of one chain with K392D/K409D (Kabat numbering) mutations while the mutIL2 was fused to the C-terminus of the second Fc-chain which harbored E356K/D399K mutations. Additionally, D265A/N297A mutations were introduced in both Fc chains to render the Fc ADCC-deficient. Expression in Expi293T cells and purification was as described above. Appropriately assembled U2S3-hFc-mutIL2 material was fractionated by SEC and the presence of individual size-appropriate polypeptides was confirmed by denaturing SDS-PAGE. A direct fusion between orthogonal ligand and mutIL2 expressed as a single polypeptide with a linker comprising glycine-serine linkages, a FLAG tag, and a 6xHis tag was also generated and purified by Ni-NTA exchange chromatography^47^. Determination of IUe activity equivalents was based on the calculation that a 4.4 μM solution of wild-type IL-2 has the equivalent of 1000 IU/μL. IL-15 with a V49D mutation, which reduced binding to IL-15Rα but retained bioactivity^24^, was similarly formatted with U2S3.

CAR-T cell proliferation in response to various cytokines or U2S3-cytokine fusions was quantified with the WST-1 Cell Proliferation Reagent (Millipore Sigma #5015944001). Briefly, CAR-T cells were pelleted and resuspended in T cell media without IL-2, dispensed into 96-well plates at 4×10^4^ cells/well, and the appropriate amount of diluted U2S3-cytokine fusions was added to achieve 30 IUe/mL or higher concentration as needed in a final assay volume of 100 μL per well. Recombinant-human IL2 and IL15 (Peprotech #200-02 and #200-15) were included as controls. After incubation for three days at 37°C, 10 μL of WST-1 was added to each well and allowed to incubate for 30-60 minutes before quantifying intensity of color development on a plate reader. Changes in the proportion of GFP^+^ CAR-expressing cells in response to U2S3-cytokine fusion were monitored by flow cytometry. To monitor activation of STAT3 or STAT5 upon cytokine-fusion engagement cells were rested overnight in TCM media without IL-2 supplementation then treated with 150 IUe/mL IL-2, IL-15, U2S3-hFc-mutIL2, or U2S3-hFc-mutIL15 for two hours before fixing and staining for intracellular phospo-STAT3 (Biolegend PE anti-STAT3 Tyr705 clone 13A3-1) and –STAT5 (BD Alexa Fluor 647 anti-STAT5 pY694 clone 47). To monitor a temporal response, treated *convertible*CAR-CD8 T cells were fixed at 0, 30, 60, and 120 minutes after exposure to cytokines or U2S3-hFc-cytokine fusions then stained.

Human PBMC stimulation and immune-phenotyping studies were performed by iQ Biosciences. Briefly normal PBMCs from three donors were seeded in 96-well plates at 1×10^5^ cells/well and exposed to a 10-fold dilution series of either U2S3-hFc-mutIL2 or U2S3-hFc-wtIL2 (wild-type IL2) for four days at 37°C with 5% CO2. Positive controls included wells coated with anti-human CD3 (OKT2) at 2 μg/mL and rhIL-2 at 300 IUe/mL. After incubation, cells were treated with TruStain FcX block (BioLegend #422301) followed by staining with BioLegend antibody panels for proliferating T cells (CD8 clone RPA-T8 #301050, CD4 clone OKT4 #317410, CD3 clone OKT3 #300430, KI-67 #350514), and Treg cells (Fox3 clone 206D #320106, CD4 clone OKT4, CD3 clone OKT3, KI-67.

## AUTHOR CONTRIBUTIONS

KEL, KCK, KTR, and DWM conceived the experiments. KCK and KEL supervised the study. SRW performed most of the *in vitro* experiments while KCK, KEL, DG, DS, and SL performed or contributed to specific experiments. KCK wrote the manuscript with editorial contributions provided by DWM, KEL, and KTR. All authors have read and approved the final submitted manuscript.

---

**Supplementary Figure 1:**
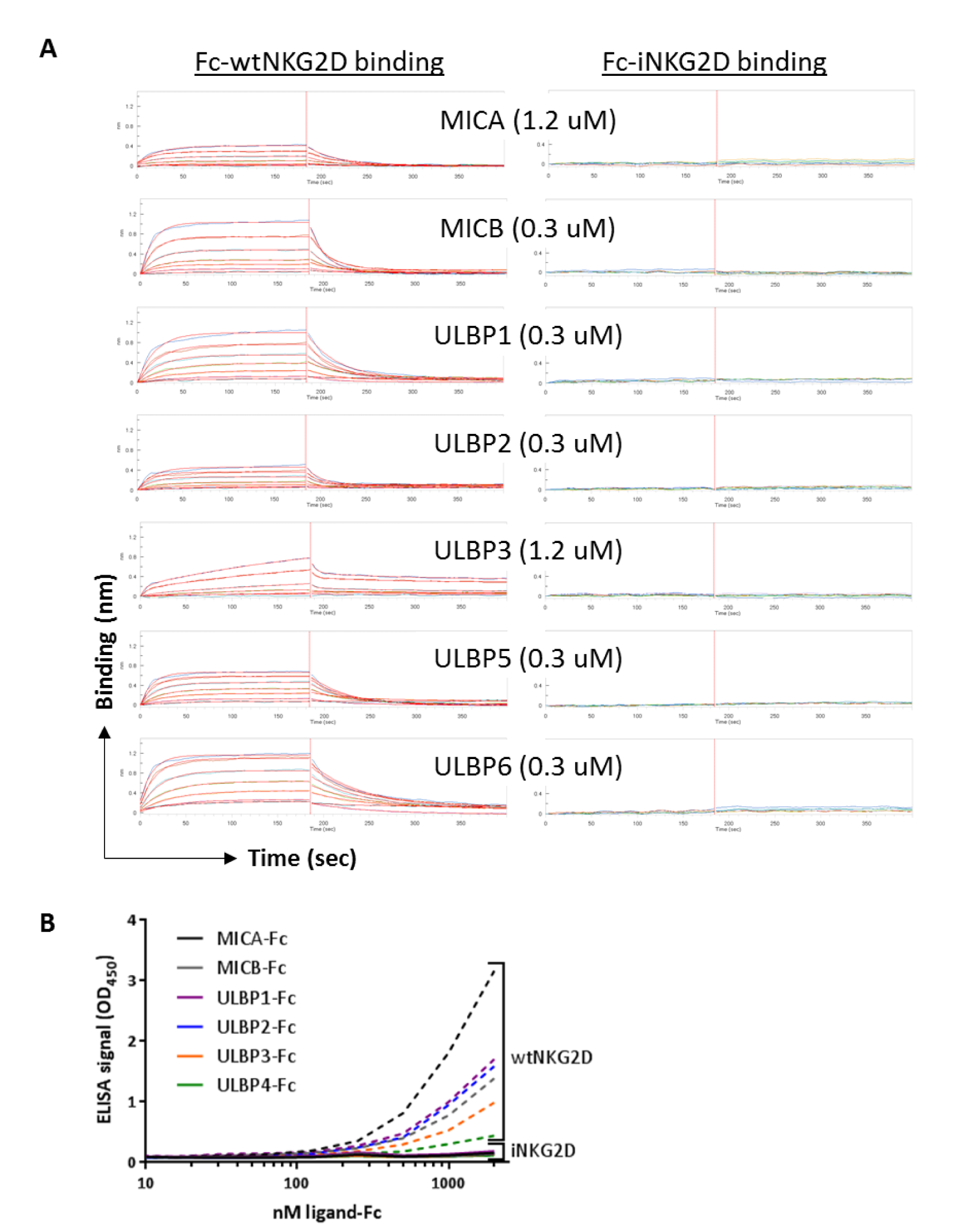
**(A)** Octet BLI kinetic binding data for His-tagged monomeric wild-type MIC ligand interaction with either wild-type NKG2D or iNKG2D.YA. Fc-wtNKG2D or Fc-iNKG2D.YA were captured with anti-human IgG Fc capture (AHC) biosensor tips associated with a dilution series of each ligand (parenthetical value indicates highest concentration examined) after baseline establishment. ULBP4 could not be expressed and purified as a monomer so was not included in this assay. Note that all axes are to the same scale. **(B)** ELISA confirming inability of iNKG2D.YA to engage natural ligands. Ligand-Fc fusions (R&D Biosystems) were coated onto microtiter plates and a titration of biotinylated Fc-wtNKG2D (dashed lines) or Fc-iNKG2D.YA (solid lines) applied and detected by streptavidin-HRP.

**Supplementary Figure 2:**
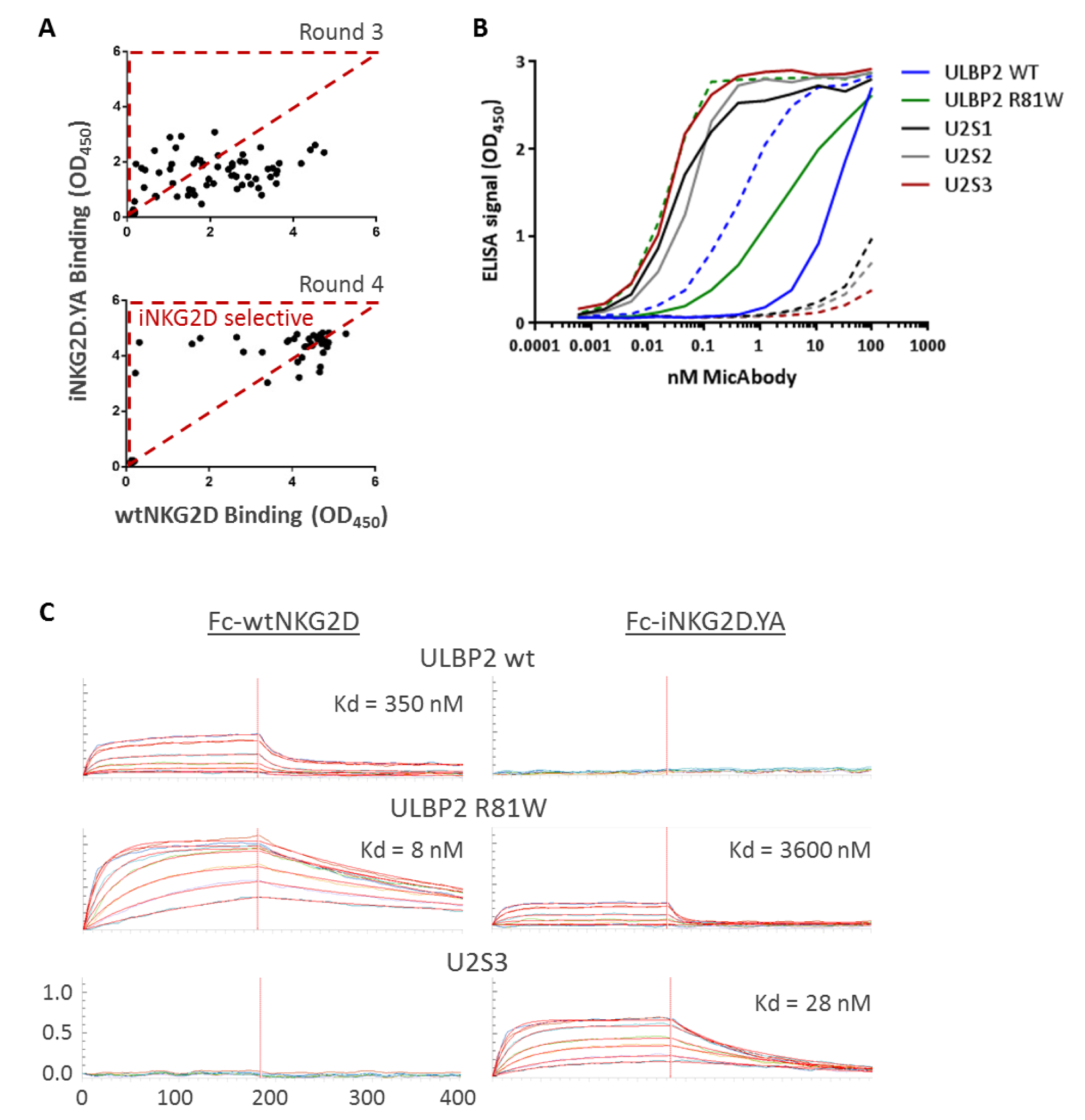
**(A)** Relative binding of selected phage to Fc-iNKG2D.YA and Fc-wtNKG2D after the third and fourth rounds of panning in the presence of increasing concentrations of wtNKG2D competitor. Phage clones in the portion of the graph outlined by the red triangle were selected for further characterization. **(B)** Three phage variants – S1, S2, S3 – were expressed as fusions to the C-terminus of the anti-FGFR3 antibody clone R3Mab heavy chain as MicAbodies and, along with wild-type ULBP2 and R81W versions, were tested for the ability of the selective variants to retain preferential Fc-iNKG2D.YA binding (solid lines) over Fc-wtNKG2D (dashed lines). All purified MicAbodies retained binding to human FGFR3 (data not shown). **(C)** Binding analysis of His-tagged monomeric wild-type ULBP2, ULBP2 R81W, and the orthogonal U2S3 ligand binding to Fc-NKG2D and Fc-iNKG2D.YA. Fc-wtNKG2D or Fc-iNKG2D.YA were captured with anti-human IgG Fc capture (AHC) biosensor tips then associated with a dilution series of ligand.

**Supplementary Figure 3:**
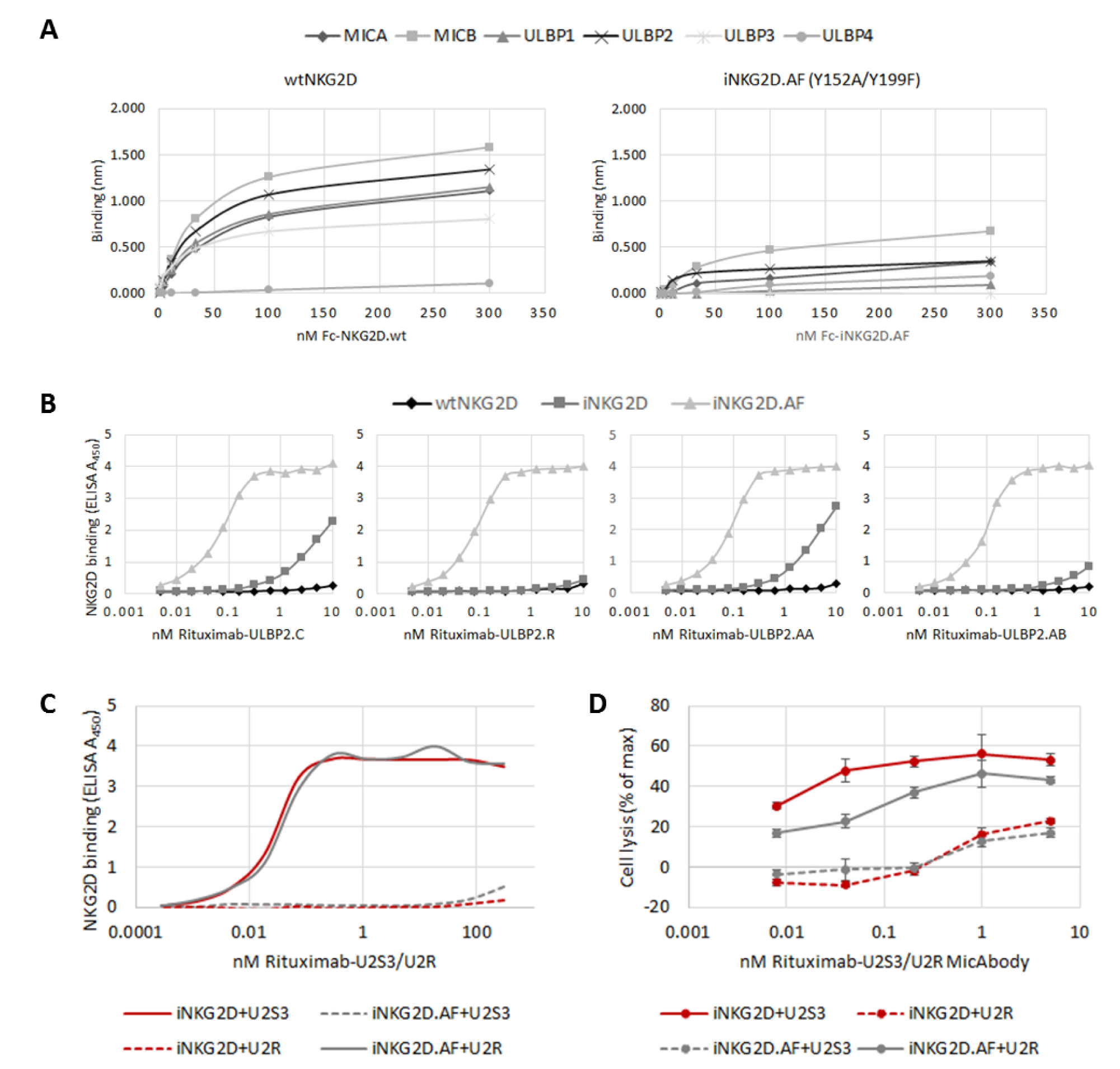
Orthogonal ULBP2 ligand selective binding to NKG2D Y152A/199F (iNKG2D.AF). Library design and phage panning performed as described for iNKG2D.YA except that biotinylated double-mutant Fc-iNKG2D.AF was used during rounds of selection against increasing concentrations of Fc-wtNKG2D competitor. **(A)** Octet BLI binding data for interaction of monomeric ligands to either Fc-wtNKG2D or Fc-iNKG2D.AF. **(B)** Lead variants selected from the phage display library were cloned as fusions to the C-terminus of the rituximab light chain and differential binding to Fc-wtNKG2D, Fc-iNKG2D.YA, and Fc-iNKG2D.AF and quantified by ELISA. Shown are four variants that selectively engage Fc-iNKG2D.YA and not the other two receptors. **(C)** ELISA demonstrating exclusivity of U2S3 and U2R ligand binding to the receptor variant against which it was selected – Fc-iNKG2D.YA and Fc-iNKG2D.AF, respectively. **(D)** Calcein release assay with Ramos target cells at an E:T of 20:1 with either iNKG2D.YA-CAR or iNKG2D.AF-CAR expressing CD8^+^ T cells and a titration of Rituximab.LC-U2S3 or Rituximab.LC-U2R.

**Supplementary Figure 4:**
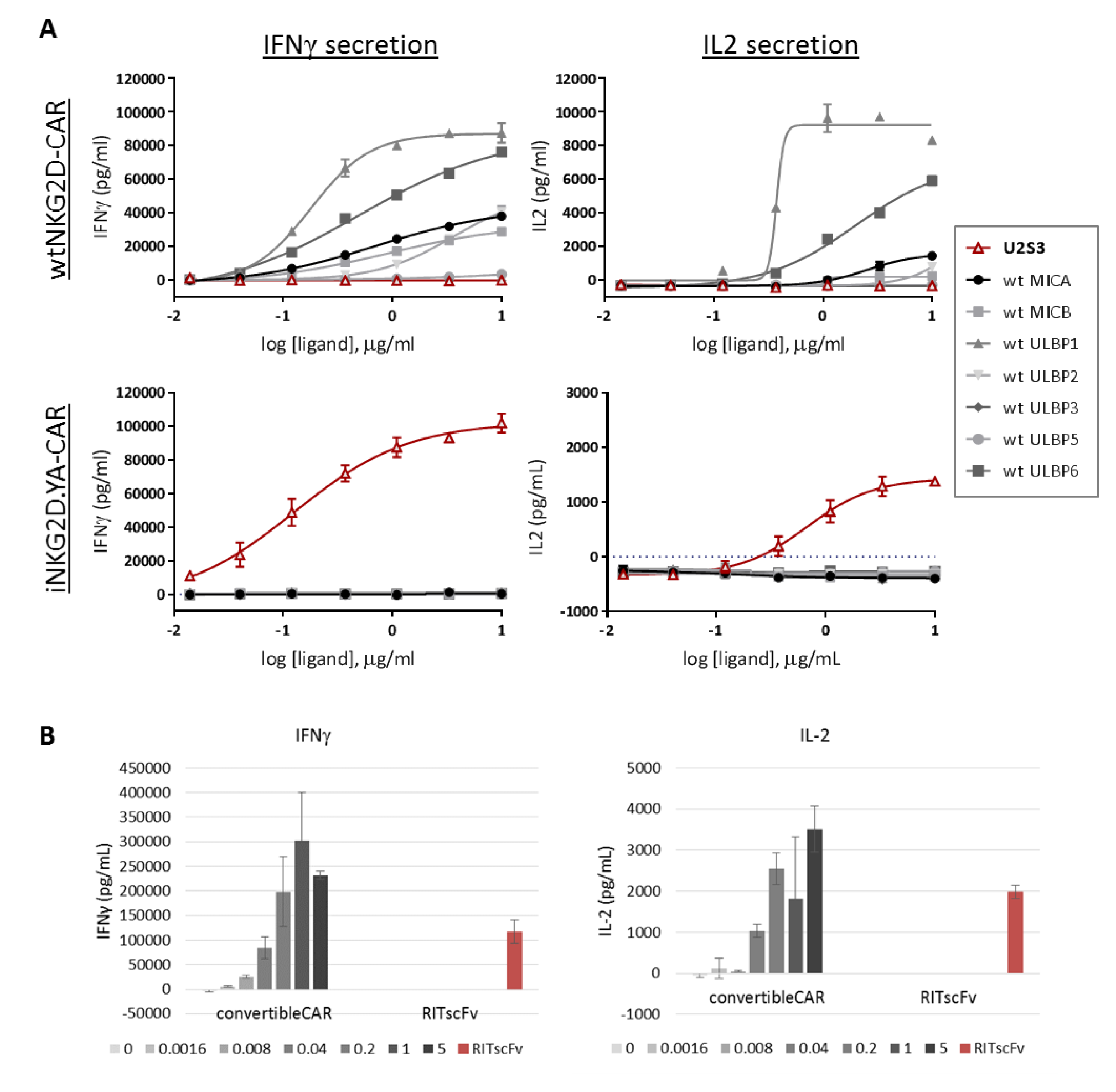
Ligand-dependent activation of iNKG2D-CAR expressing CD8^+^ T cells. **(A)** CD8^+^ T cells were transduced with CAR constructs comprised of either wild-type NKG2D or iNKG2D.YA as the receptor domain. Wild-type His-tagged monomeric ligands or His-tagged monomeric U2S3 were coated onto the wells of a microtiter plate in a 1:3 dilution series starting at 10 μg/mL. 1×10^5^ CAR expressing cells were introduced to the wells in 150 μL volume without exogenous IL2, supernatants collected 24 hours later, and the amount of cytokine produced and release quantified by cytokine-specific ELISA. ULBP4 was not included in the assay as a His-tagged version could not be expressed and purified. **(B)** CD8^+^ cells expressing either iNKG2D-CAR or RITscFv-CAR were co-cultured with Ramos cells at an E:T of 4:1 with increasing concentrations of Ritux-S3 MicAbody in the case of iNKG2D-CAR cells. After 24 hours, culture supernatants were harvested and released cytokine quantified by ELISA. All error bars are ±SD of technical triplicates.

**Supplementary Figure 5:**
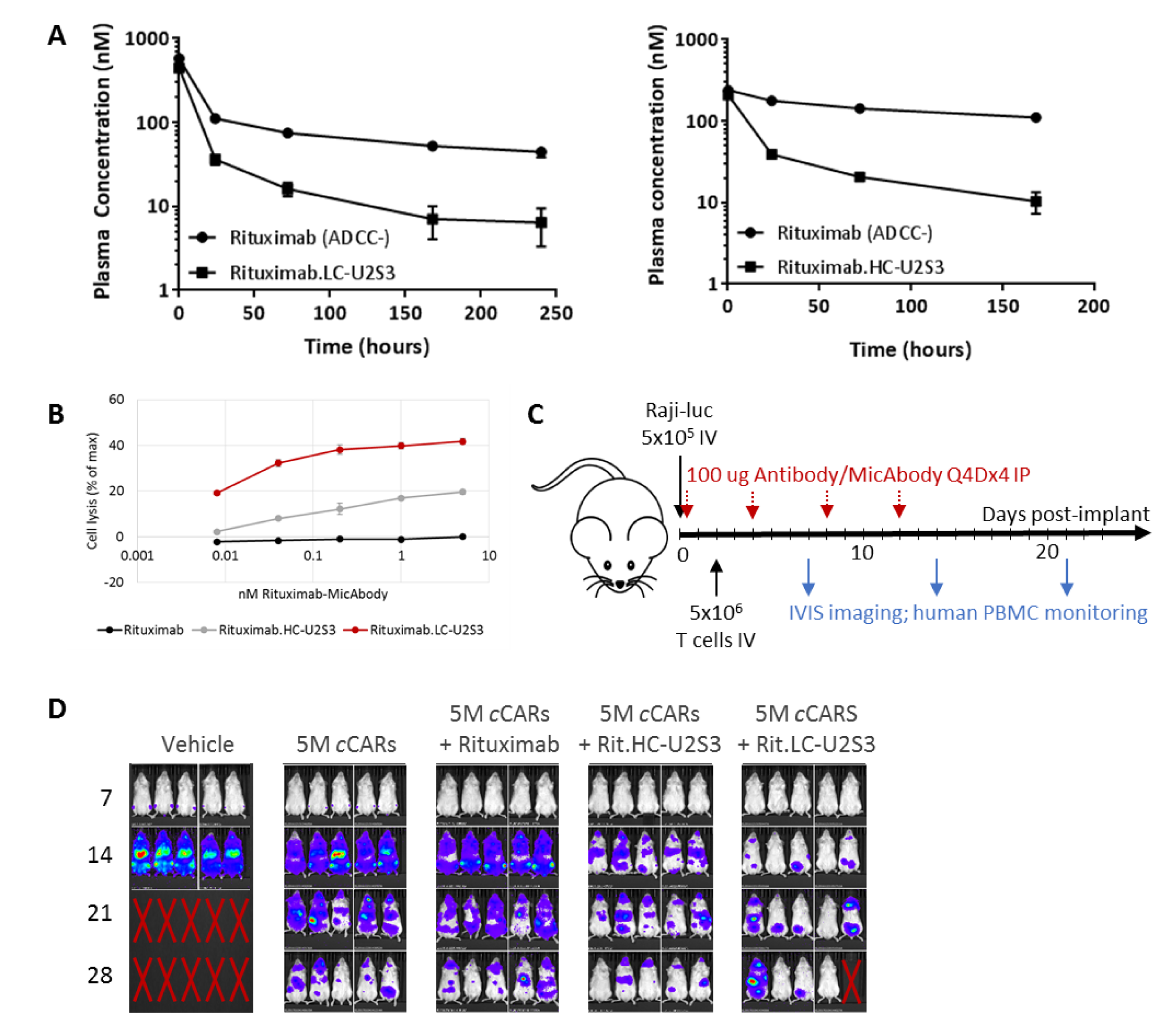
Comparison of heavy-vs. light-chain U2S3 fusions to rituximab (ADCC-deficient) antibody. **(A)** Pharmacokinetics of serum Rituximab-U2S3 MicAbody levels after 100 μg IV administration in NSG mice in the absence of human T cells or tumor. Note that all MicAbodies and antibody controls used were ADCC-deficient. The graph on the left is a comparison of parental antibody to the light-chain U2S3 fusion while the graph on the right is a comparison of parental antibody to the heavy-chain U2S3 fusion. All error bars are ±SD of technical triplicates. **(B)** *In vitro* calcein release assay after two hours co-culture with iNKG2D-CAR CD8^+^ T cells and Ramos target cells at an E:T of 20:1 and titrations of Rituximab-MicAbodies. Error bars represent ±SD for the experiment and data are representative of multiple experiments. **(C)** *In vivo* study design in NSG mice with Raji-luciferase cells invused IV followed by treatment and monitoring as indicated. CD4^+^ and CD8^+^ cells were independently transduced, combined at a 1:1 ratio without adjusting for percent transduction, and a total of 5×10^6^ *convertible*CAR-T cells (*c*CARs) were injected IV. **(D)** IVIS imaging was performed 7, 14, 21, and 28 days post-implantation and all adjusted to the same scale. Death of mouse #5 at day 28 of the 5M iNKG2D + Rituximab.LC-U2S3 was unrelated to treatment or disease.

**Supplementary Figure 6:**
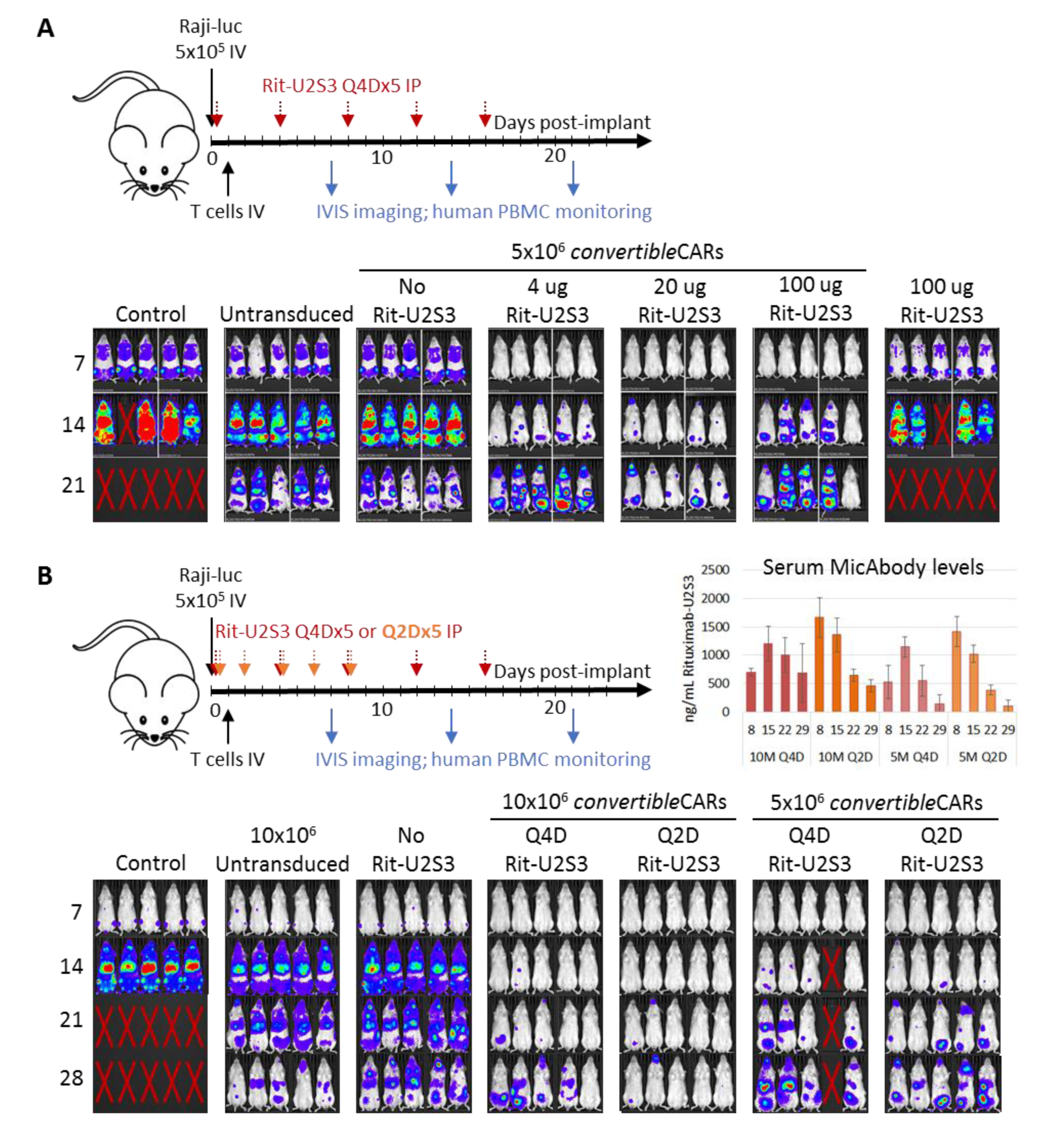
Investigation of both MicAbody and *convertible*CAR dosing strategy in a disseminated Raji-luciferase B cell lymphoma NSG mouse model. **(A)** Rituximab.LC-U2S3 Q4Dx5 dosing was initiated the same day as tumor implantation at 4, 20, or 100 μg per dose while keeping the number of cells administered constant across cohorts. The 100 μg Rituximab.LC-U2S3 only cohort – without any CAR-T cells – received just a single dose of MicAbody. **(B)** Frequency of MicAbody dosing as well as *convertible*CAR-T cell infusions levels were explored with the former being administered either every two or four days for a total of 5 doses, and the latter at either 5×10^6^ or 10×10^6^ total cells infused. Mouse #4 in the Q4D + 5×10^6^ cohort that died by day 14 did so for reasons unrelated to treatment or disease. Serum levels of Rituximab.LC-U2S3 were also monitored at 8, 15, 22, and 29 days post-implantation for all cohorts that received MicAbody (cohorts that did not receive any were negative by ELISA, data not shown). Error bars are ±SD of triplicate measurements. A non-CAR specific graft-vs-tumor effect was clearly observed in this study with the untransduced and no MicAbody cohorts. IVIS images within each respective study were adjusted to the same scale.

**Supplementary Figure 7:**
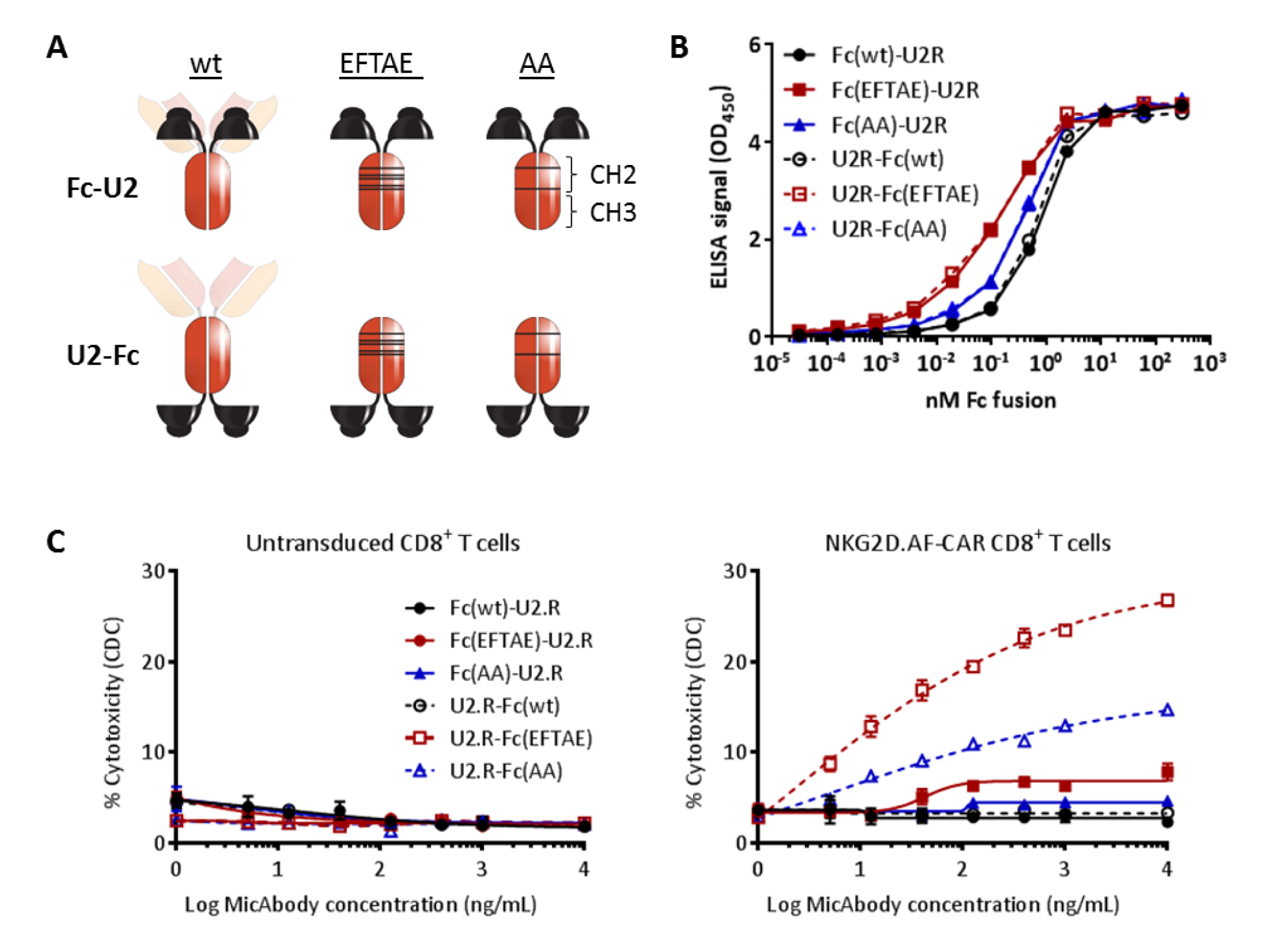
Targeted recruitment of complement factor C1q to iNKG2D.AF-CAR cells to direct their complement-mediated attrition. **(A)** Structure of orthogonal ligand fusions to the Fc portion of human IgG expressed as either N- or C-terminal fusions. In addition to wild-type Fc, two sets of mutations in the CH2 domain that enhance C1q binding were independently explored - S267E/H268F/S324T/G236A/I332E (“EFTAE”) and K326A/E333A (“AA”). **(B)** ELISA examining binding of human C1q to each purified fusion protein. Rank order of Kd’s was EFTAE<AA<wt (0.12, 0.35, and 0.67 nM, respectively) regardless of orientation of fusions. **(C)** Complement-dependent cytotoxicity (CDC) assays for C1q-binding enhance Fc-fusions. iNKG2D.AF-CAR (59% GFP^+^) or untransduced CD8^+^ T cells were incubated with a titration of each fusion molecule and 10% normal human serum complement for three hours before dead T cells were enumerated with SYTOX Red. Similar results were obtained with U2S3 orthogonal ligand fusions to direct complement-enhanced Fc domains to iNKG2D.YA-CAR cells (data not shown). All error bars are ±SD of triplicate measurements.

**Supplementary Figure 8:**
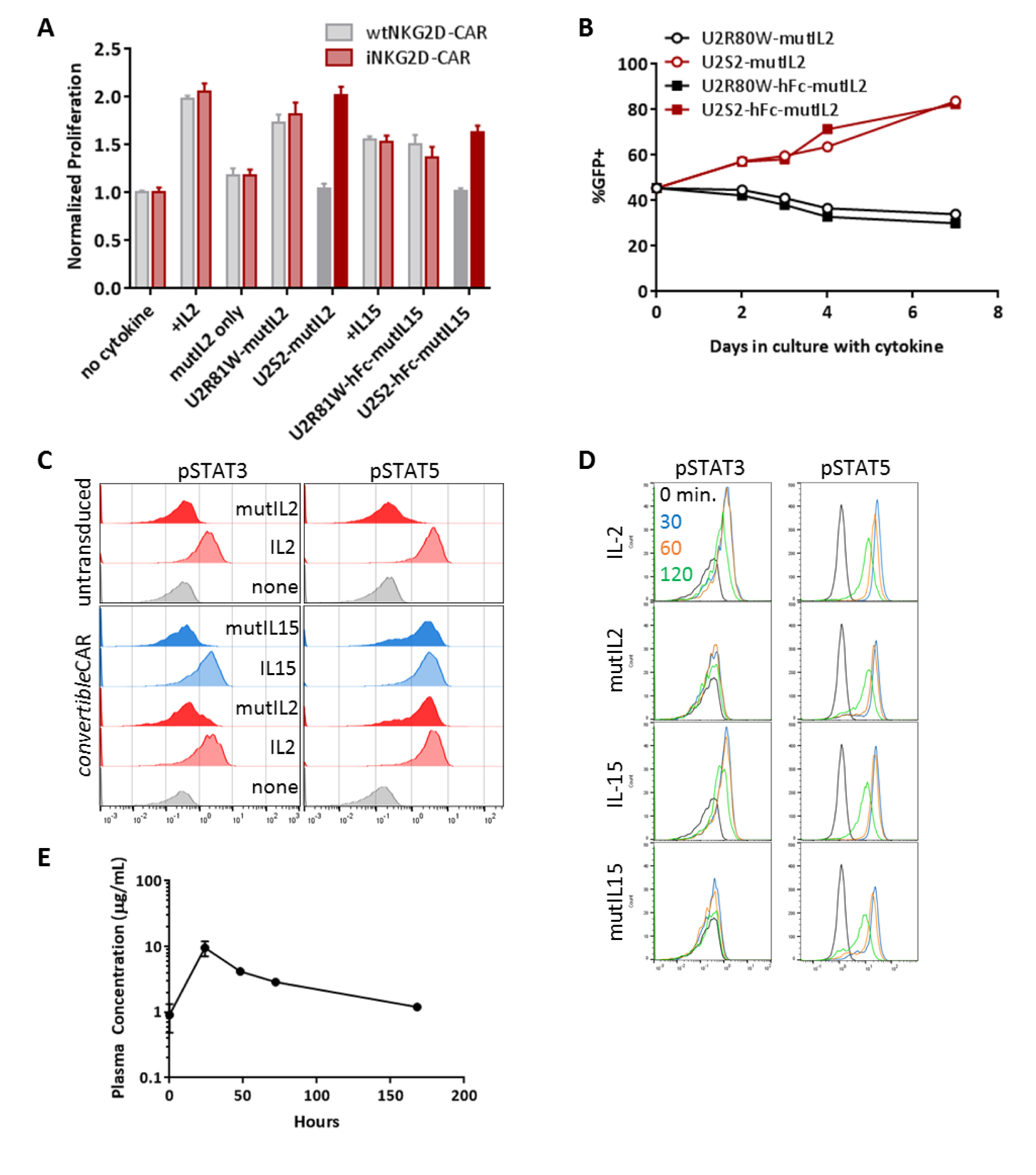
Targeted delivery of mutant-IL2 cytokine to iNKG2D.YA-CD8^+^ T cells. **(A)** *in vitro* proliferation after three days of wtNKG2D-CAR or iNKG2D.YA-CAR treatment with 30 IUe/mL of cytokine or cytokine-U2S2 fusion. Darker shading is to highlight selectivity. **(B)** A low efficiency (45% GFP^+^) iNKG2D.YA-CAR transduction was cultured with 30 IUe/mL of non-selective (U2R81W) or iNKG2D.YA-selective (U2S2) mutIL2 fusion and maintained for seven days. Cells were periodically examined by flow cytometry to quantify the %GFP^+^ cells in each population. **(C)** Untransduced or *convertible*CAR-CD8 T cells were starved overnight of supporting cytokine then treated with 150 IUe/mL IL-2, IL-15, U2S3-hFc-mutIL2, or U2S3-hFc-mutIL15 for two hours before fixing and staining for intracellular phospo-STAT3 and –STAT5. **(D)** *convertible*CAR-CD8^+^ T cells were treated as in (C) except that cells were fixed at 0, 30, 60, and 120 minutes after exposure to cytokines or U2S3-hFc-cytokine fusions then stained for intracellular phospo-STAT3 and –STAT5. **(E)** Serum PK of U2S3-hFc-mutIL2 after 60 μg IP injection in NSG mice. All error bars are ±SD of technical triplicates.

**Supplementary Figure 9:**
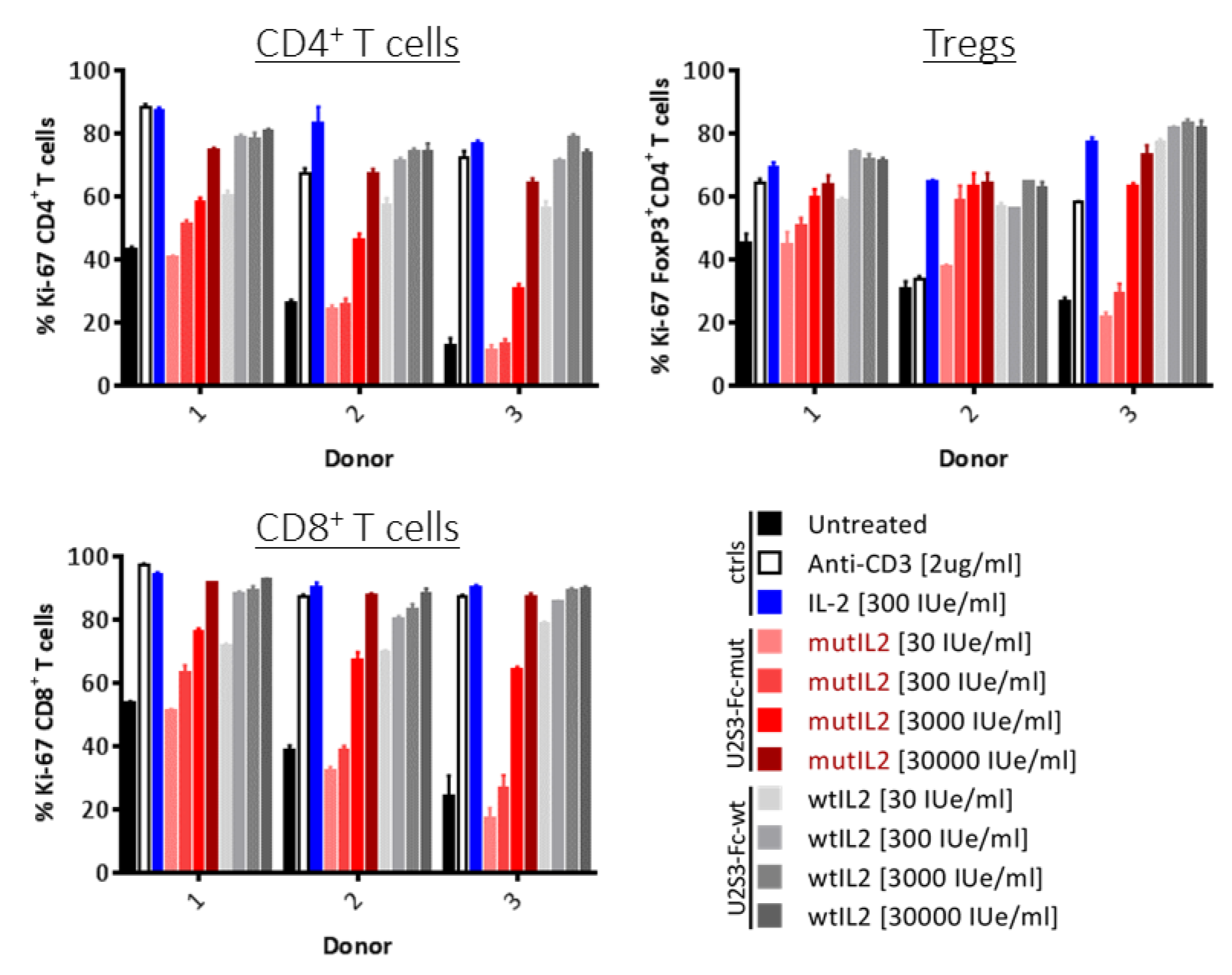
Responsiveness of human PBMCs to U2S3-hFc-mutIL2. Human PBMCs from three donors were incubated with increasing concentrations of U2S3-hFc-mutIL2 or U2S3-hFc-wtIL2 for four days along with controls. Each of the labeled cell types was examined for the marker Ki-67 to quantify proliferative response under each condition. Error bars are ±SD of triplicate measurements.

